# Altered lysosomal biology impairs motor neuron survival via TFEB dysregulation in spinal muscular atrophy

**DOI:** 10.1101/2025.07.07.663525

**Authors:** Ines Rosignol, Zeynep Dokuzluoglu, Antonio Caldarelli, Ana-Maria Oprişoreanu, Sofiia Ushakova, Tohid Siddiqui, Rahul Grover, Hannes Guerlich, Björn Falkenburger, Stefan Diez, Tobias Grass, Catherina G. Becker, Natalia Rodriguez-Muela

**Affiliations:** German Center for Neurodegenerative Diseases e.V. (DZNE), Dresden, Germany; Technische Universität Dresden, Center for Regenerative Therapies Dresden (CRTD), Dresden, Germany; B CUBE - Center for Molecular Bioengineering, TU Dresden University of Technology, Dresden, Germany; Department of Neurology, University Hospital Carl Gustav Dresden, Dresden, Germany; Max Planck Institute for Molecular Cell Biology and Genetics, Dresden, Germany; Centre for Discovery Brain Sciences, University of Edinburgh Medical School, Edinburgh, UK; Cluster of Excellence Physics of Life, TU Dresden, Dresden, Germany

## Abstract

Spinal muscular atrophy (SMA) is a devastating motor neuron disease, caused by recessive mutations or deletions of the *SMN1* gene, representing the leading genetic cause of infant mortality. Available therapies, aimed at increasing SMN protein levels, can only partially halt motor neuron (MN) degeneration in a select number of patients, reinforcing the need for combinatorial treatments to improve clinical outcomes. We previously showed that mTORC1 overactivation and impaired autophagosome clearance in SMA MNs lead to the accumulation of protein aggregates, contributing to MN degeneration. However, the mechanistic link between SMN protein deficiency and autophagy-lysosomal dysfunction remained unknown. Here, using patient iPSC-derived MNs along with isogenic and healthy controls, we show that SMA MNs exhibit reduced lysosome numbers and impaired functionality. Furthermore, the master regulator of lysosomal biogenesis and autophagy, TFEB, is downregulated, and its nuclear translocation compromised upon SMN deficiency. We further propose the upregulation of the mTORC1 positive modulator TPT1 as contributor to TFEB dysregulation. Notably, TFEB overexpression ameliorates protein aggregate accumulation in SMA MNs and enhances MN survival both in vitro and in a zebrafish SMA model. Our findings identify lysosomal dysfunction as a key player in SMA pathology and highlight TFEB activation as a potential therapeutic strategy for SMA treatment.

**One Sentence Summary:** TFEB activation restores lysosomal function and improves motor neuron survival in SMA, highlighting its potential as a therapeutic target.

## Introduction

Spinal muscular atrophy (SMA) is a neuromuscular disease and the leading monogenetic cause of infant mortality (*1, 2*). It is caused by homozygous mutations or deletions in the *SMN1* gene, which encodes the survival of motor neuron (SMN) protein. The resulting reduction in SMN protein levels leads to motor neuron (MN) degeneration in the ventral horn of the spinal cord, severe muscle weakness, and often premature death (*1*). In 1995, researchers identified the genetic cause of SMA, which is the *SMN1* gene (telomeric) and its paralogue *SMN2* (centromeric), located to chromosome 5q11.1-13.3 (*3*). This discovery was pivotal in understanding SMA pathogenesis. Both *SMN* genes are constitutively transcriptionally active and almost identical, with a critical difference within exon 7 in a single translationally silent coding variant, c.840C>T. This small but significant change alters the splicing of *SMN2* transcripts (*4, 5*). The protein produced from those transcripts, termed SMNΔ7, is truncated and quickly degraded. Importantly, exon 7 is retained in about 10% of *SMN2* transcripts which therefore produce small amounts of full length, functional SMN protein (*6*). Because of the small production of full length SMN, *SMN2* is the main SMA disease modifier. Humans naturally have between 0 to 5 copies of the *SMN2* gene (*1*). Minor full length SMN production from *SMN2* makes *SMN2* copy number a critical determinant for disease severity, wherein a higher copy number equals higher full length SMN amount and results in a less severe presentation of SMA. Full-length SMN protein is ubiquitously expressed and essential for the survival of organisms across the animal kingdom (*7, 8*). Its critical role underscores its importance in maintaining cellular function and organismal viability. Even though SMN has been linked to many cellular homeostasis pathways, the first role of SMN discovered, and best characterized, is its chaperone function in the assembly of the spliceosome by aiding the biogenesis of small nuclear ribonucleoproteins (snRNPs) (*9, 10*). More roles of SMN have been described in cellular housekeeping processes, such as mRNA transport, protein translation, cytoskeletal dynamics, bioenergetics pathways, stress granule formation, endocytosis and protein homeostasis pathways, including the UPS and autophagy (*11*), but some of these are mechanistically less understood.

Alterations in autophagy are associated with a wide range of adult-onset neurodegenerative diseases (NDs) and certain neurodevelopmental disorders, including autism spectrum disorder, microcephaly, Vici syndrome, hereditary spastic paraplegia and Charcot-Marie-Tooth (*12*). In fact, the buildup of neurotoxic protein aggregates in the cytosol is a hallmark shared by numerous NDs. Notably, individuals with certain hereditary neurodegenerative and neurodevelopmental conditions often carry loss-of-function mutations or polymorphisms in core autophagy-related genes (ATGs) or genes encoding proteins crucial for autophagosome (AP) trafficking, maturation, and lysosomal fusion (*12*). Additionally, the early stages of sporadic NDs are often marked by the convergence of age-associated neuroinflammation and disruptions in autophagy (*13*). However, the precise connections between autophagy and the development of NDs have yet to be fully clarified. Even though a dysregulation of the autophagy-lysosome pathway (ALP) has been proven a critical factor in almost all NDs, including other motor neuron diseases (MNDs), such as amyotrophic lateral sclerosis (ALS) or Charcot-Marie-Tooth, the role it plays in SMA has been neglected. The common ground of the few available studies is the accumulation of APs accompanied by increased levels of autophagy cargo and selective adaptor p62 and a defective autophagy flux (*14–17*). Additionally, we and others reported an abnormal overactivation of the mammalian target of rapamycin complex 1 (mTORC1) as a potential cause of these ALP alterations in SMA (*14, 18*), however, the molecular underpinnings of the autophagy dysregulation and how exactly SMN regulates this process are still unanswered questions.

In this study, we used type 2 and type 1 SMA hiPSC lines, along with their recently generated genome-engineered isogenic controls (*19*) and an additional healthy hiPSC line, to investigate how ALP dysregulations contribute to SMA pathophysiology. Our findings reveal that MNs derived from the SMA hiPSC lines exhibit reduced numbers of lysosomes and autolysosomes, accompanied by diminished autophagy-lysosome-mediated proteostasis capacity and clear signs of intracellular protein aggregation. We found that these defects are likely due to a dysregulation of TFEB nuclear translocation originated from increased TPT1-mTORC1 activity levels. These phenomena seemed specific to MNs as no similar decrease in survival or appearance of lysosomal defects were observed in SMA cortical neurons differentiated from the same patient-derived hiPSCs and could at least partially explain the observed selective vulnerability of spinal MNs in SMA patients. Importantly, overexpression of TFEB in vitro and in a zebrafish model of SMA rescued the protein aggregation phenotype and MN death. This study highlights the importance of this catabolic axis in the pathophysiology of SMA MNs, and thus suggests that boosting lysosomal/autophagy functionality could constitute a promising new strategy to use in combination with the current SMN-increasing therapies.

## Results

### SMA patient hiPSC-derived motor neurons have less lysosomes than the isogenic control and healthy motor neurons

Previous studies have reported that SMN-deficient cells show defects in the endosomal-lysosomal system (*17, 20*) and an accumulation of autophagy substrates, ubiquitinated proteins and undegraded autophagosomes (APs) (*14, 15*). However, the mechanism connecting SMN loss to these alterations in the ALP is still unknown. The fact that APs still form properly but accumulate in SMN-deficient cells (*14*) suggests a failure in the final step of the autophagy flux, specifically in the lysosome-mediated degradation of APs. Therefore, we explored whether an SMN reduction could lead to decreased lysosome biogenesis, potentially causing the aberrant accumulation of APs. First, we used a HEK293T cell line stably expressing a doxycycline inducible shRNA against SMN and observed that SMN knockdown (KD) resulted in a significant decrease in the amount of lysosomal-associated membrane protein 1 LAMP1 and the lysosomal hydrolase cathepsin B (CTSB) at the proteins levels (Figure S1A-E), which suggested a reduced number of lysosomes in the SMN-depleted cells. Next, we explored whether a SMN reduction caused similar lysosomal changes in human MNs. We applied a well-established protocol to differentiate hiPSCs into spinal MNs following specific developmental programs (*14, 19, 21*) (Figure 1A) on a SMA hiPSC isogenic model that we recently developed, composed of a mild SMA type 2 line, a severe SMA type 1 line and one isogenic healthy clone for each line generated by genome-editing mediated conversion of one *SMN2* gene into *SMN1* (*19*). As additional control to help determine a non-disease baseline, a healthy hiPSC line (WT) reported in numerous studies before, was used (*14, 19, 21–29*). First, to confirm that our model recapitulated the major SMA disease hallmark, increased MN death over healthy controls, we measured the survival of these hiPSC-derived MNs via longitudinal time-laps over a 10-days period and observed an initial phase of acute cell death (first 4 days in vitro post EB dissociation and MN plating), followed by a second phase of more steadily decline in survival (Figure 1B-C). These results align with the bi-phasic MN degeneration profile that has been described in patients affected by severe forms of SMA (*30*). MN survival over time matched the severity of the disease lines and was significantly ameliorated in the isogenic controls (Figure 1B-C). Interestingly, unbiased automated single-cell quantification of SMN protein levels in the individual MNs at an early (DIV2) and late (DIV10) time-point in culture revealed that the long-living MNs harboured significantly higher SMN levels than the ones present in the early cultures, independent of their disease or healthy genotype (Figure 1D and S2). In addition, DIV10 SMA type 2 and 1 MNs showed on average similar SMN protein levels to DIV2 healthy MNs (Figure 1D). These results confirm our previous study on the heterogeneity of SMN protein levels that individual MNs express and how this inter-MN variability determines their probability of survival (*21*). These data suggest that healthy and SMA early MN cultures are composed by a heterogeneous mixture of high- (“resistant”) and low-SMN (“vulnerable”) pools of MNs and the late cultures primarily contain the surviving high-SMN MNs.

**Figure 1.**
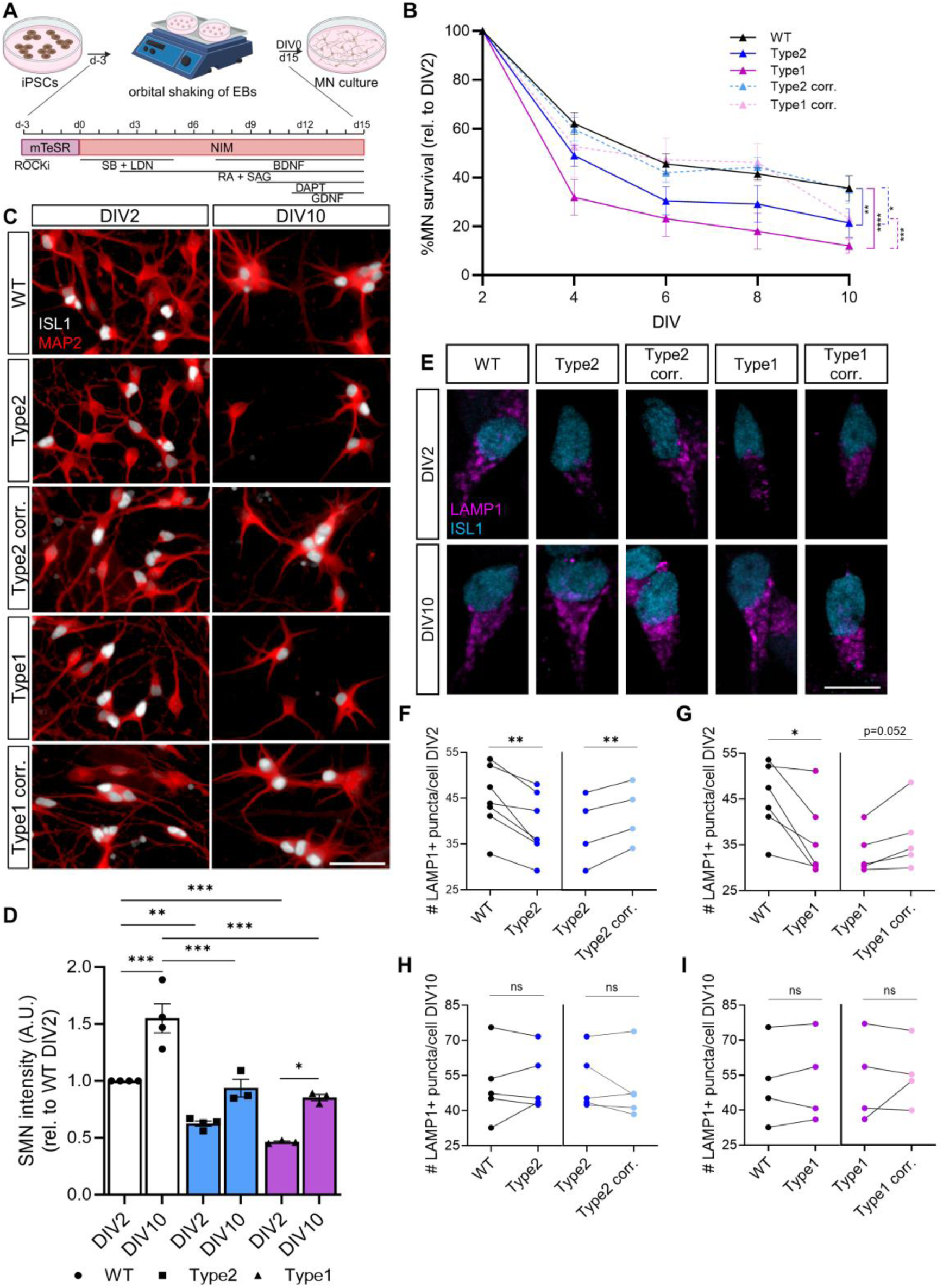
Reduction of SMN protein levels leads to a decreased number of lysosomes. (**A**) Schematic representation of protocol followed for MN generation from hiPSCs. (**B**) Quantification of SiR-DNA-labeled WT, SMA type 2 and type 1 MN survival over time (results are shown relative to DIV2 for each line, N= 3-6 individual experiments, each representing the mean of n= 3 technical replicates per line per time-point; two-way ANOVA with Tukey’s multiple comparison test). (**C**) Representative images of hiPSC-derived MN cultures immunostained against ISL1 (white) and MAP2 (red) 2 and 10 days after plating post-EB dissociation. Scale bar, 50µm. (**D**) Quantification of immunostained SMN protein levels in WT and SMA MNs over time (Each independent experiment is plotted as a circle, each representing the mean of n= 3 technical replicates; two-way ANOVA with Tukey’s multiple comparison test). (**E**) Representative images of DIV2 and DIV10 hiPSC-derived MNs immunostained against LAMP1 (magenta) and ISL1 (blue). Scale bar, 10µm. (**F-I**) Quantification of the number of LAMP1^+^ puncta per MN at DIV2 in SMA type 2 vs isogenic control and WT MNs (**F**), SMA type 1 vs isogenic control and WT MNs (**G**) and similar analysis from DIV10 MNs (**H-I**). Each independent experiment is plotted as individual circle, each representing the mean of n= 3 technical replicates. Samples from the same experiment are connected by a line. 2-tailed Paired t-test. Mean ± SEM is shown.

To determine if lysosomal numbers were altered in SMA MNs, quantification of the amount of LAMP1^+^ and CTSB^+^ puncta was performed in DIV2 and DIV10 ISL1^+^ MNs. A significant decrease in the number of LAMP1^+^ lysosomes in both DIV2 SMA type 2 and type 1 MNs compared to their respective isogenic controls and to the WT was observed (Figure 1E-G), and, interestingly, no longer detected in the DIV10-surviving MNs (Figure 1E, H-I). The analysis of the number of CTSB^+^ puncta per ISL1^+^ MNs confirmed these findings (Figure S3A-E). We also quantified individual lysosome size and observed that lysosomes are smaller in SMA type 2 and 1 MNs than in the controls, specifically in the early cultures composed by vulnerable and resistant MNs (Figure S3F-I). Taken together, these data shows that the most vulnerable SMA MNs, dying faster in culture, contain a reduced number of lysosomes and lysosomal hydrolases than the isogenic and healthy controls and the more resilient SMA MNs. These findings thus support a previously unknown function of SMN in the regulation of lysosomal biology.

### Lysosomes in SMA MNs show an increased acidity and CTSB activity

The lysosome is a critical organelle for maintaining protein homeostasis by degrading cellular components via proteases that reside within its lumen (*31*). Lysosomes have a characteristic acidic pH of 4-5, which is essential for protease activity, fusion with autophagosomes and the effective clearance of macromolecules and whole organelles, such as mitochondria (*32*). Lysosomal pH is maintained by v-ATPases, transmembrane proton pumps localized in lysosomes, endosomes and secretory vesicles (*33*). A previously published RNAseq dataset (*23*) revealed that several subunits of the v-ATPase were increased in SMA type 1 MNs compared to WT MNs (ATP6V1A, ATP6V1E1, ATP6V0E1, ATP6V1C1 and ATP6V1G1). Furthermore, ATP6V0E1 and ATP6V1G1 also appeared significantly upregulated in a single-cell RNAseq study on neuromuscular organoids developed from SMA versus isogenic control and healthy hiPSCs that we recently published (Figure S3J) (*19*). Since pH is a key parameter for the proper functioning of lysosomes, we questioned whether the previously observed increase in v-ATPase subunits in SMA MNs translated to an abnormal lysosomal acidity. DIV2 and DIV10 MN cultures were stained with the live dyes LysoSensor or LysoTracker (LTR) to label acidic vesicles. LysoSensor, an acidotropic molecule that accumulates in a broad range of acidic vesicles with pH 4.5-6 (*34*), was used in combination with the live dye CalceinRed to stain the somas of living cells. Automated image-based analysis of the fluorescence intensity distribution of all LysoSensor^+^ puncta per cell revealed that the severe SMA type 1 MNs show an increased acidic level of vesicles compared to healthy controls, both at DIV2 and DIV10 (Figure 2A-C). As the LysoSensor dye exhibits a pH-dependent increase in fluorescence signal, this suggests that, in agreement with the mentioned RNAseq study, SMA type 1 MNs present increased vesicle acidification compared to type 2 and control MNs.

**Figure 2.**
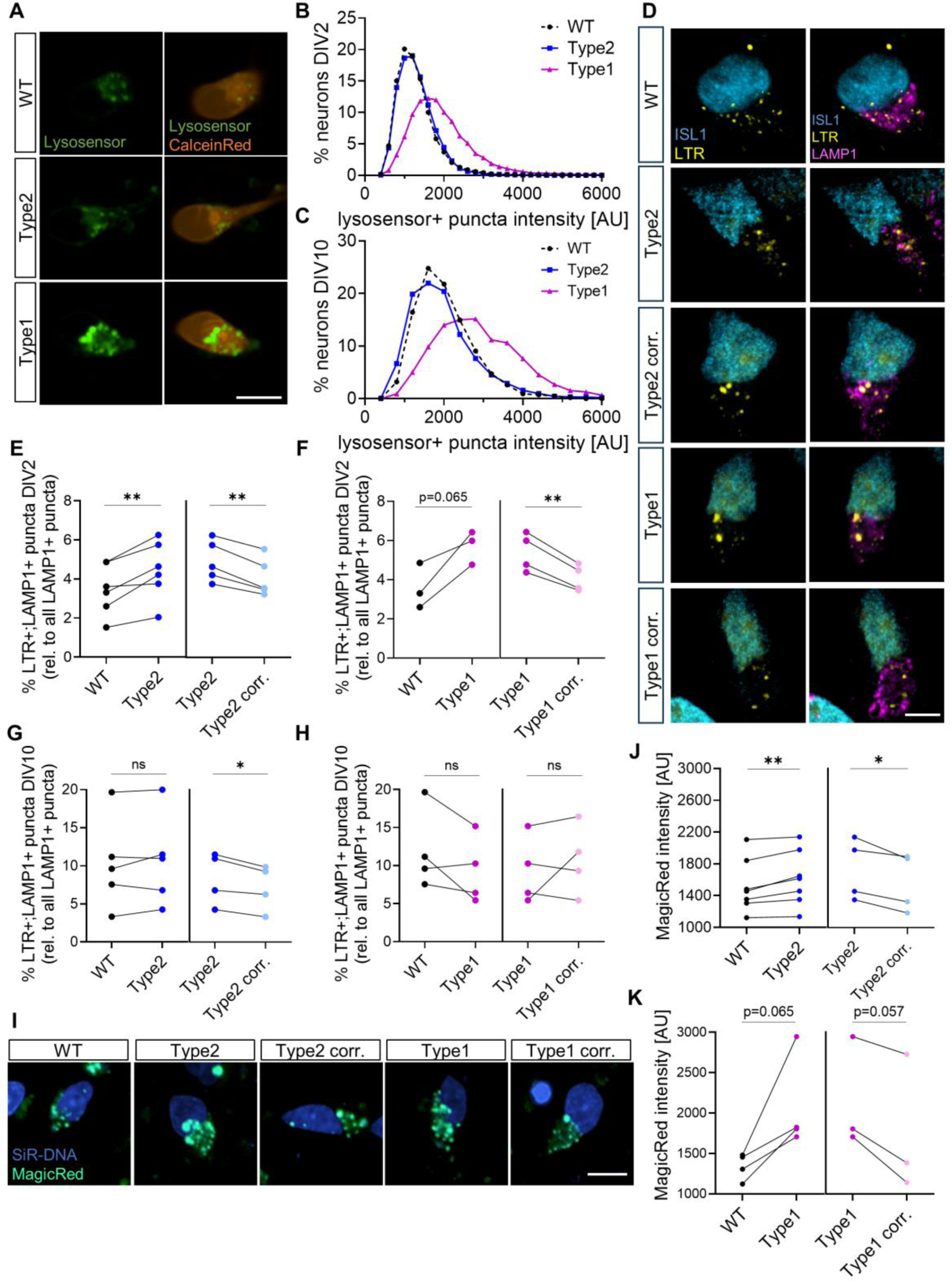
Lysosomes in SMA MNs show an increased acidity and CTSB activity specifically in early cultures (composed by vulnerable and resistant MNs). (**A**) Representative images of DIV2 WT and SMA MN cultures labeled with live pH probe lysosensor (green) and live cell dye calcein red (orange). Scale bar, 10µm. (**B**) Representative histogram quantification of the fluorescence intensity of each lysosensor^+^ puncta per neuron in DIV2 and (**C**) DIV10 MN cultures (N= 5 individual experiments, each representing the mean of n= 3 technical replicates). (**D**) Representative images of DIV2 SMA vs isogenic control and healthy MNs dyed with LTR (yellow) and immunostained against LAMP1 (magenta) and ISL1 (cyan). Scale bar, 10µm. (**E**) Quantification of the percentage of LTR^+^;LAMP1^+^ puncta per MN relative to the total number of LAMP1^+^ puncta at DIV2 for SMA type 2 vs isogenic control and healthy MNs, and (**F**) for SMA type 1 vs isogenic control and healthy MNs. (**G**-**H**) Similar quantifications as for E-F in DIV10 MNs. (**I**) Representative images of SMA types 2 and 1 vs isogenic controls and healthy DIV2 MNs stained with the CTSB activity live dye MagicRed (green) and the live nuclear dye SiR-DNA (blue). Scale bar, 10µm. (**J**) Quantification of the intensity of MagicRed^+^ puncta in DIV2 SMA type 2 vs isogenic and (**K**) healthy and SMA type 1 vs isogenic and healthy MNs. 2-tailed Paired t-test. Mean ± SEM is shown. For panels E-F, G-H, J-K, each circle represents an independent experiment, and each represents the mean of n= 3 technical replicates. Samples from the same experiment are connected by a line.

LysoSensor accumulates in a large variety of acidic vesicles, with a pH ranging from 4.5-6, including not only lysosomes but also amphisomes, late endosomes and various transport vesicles involved in the lysosomal maturation process (*35*). To confirm that the enhanced acidic pH values detected in SMA type 1 MNs with Lysosensor was specific to lysosomes, we next used LTR, which stains vesicles with a pH of 4.5-5, in combination with LAMP1 immunostaining. Quantification of the number of LTR^+^;LAMP1^+^ puncta per individual MN revealed that both SMA type 2 and type 1 MNs displayed a significant increase compared to their isogenic control pair and the WT MNs at the early time-point studied (Figure 2D-F). This increase in acidic lysosomes was no longer present at the late time-point (Figure 2G-H). We next sought to examine whether the observed increased lysosomal acidification in the most vulnerable SMA MNs would translate into higher lysosomal degradative capacity, as an enhanced acidity is often associated with higher protease activity (*36*). We utilized the MagicRed assay to measure CTSB activity (*37*) in living cultures and observed that SMA type 2 MNs presented a mild but significant increase in CTSB activity compared to isogenic and WT controls, which was more marked in the type 1 MN cohort (Figure 2I-K). CTSB is regulated by CTSD (*38*) and, interestingly, CTSD has been recently identified as a biomarker in SMA, as it was also reported for Alzheimer’s (AD) (*39*) and ALS (*40*). CTSD levels declined in the cerebrospinal fluid of SMA patients “responders” to nusinersen treatment compared to “non-responders” (*41*). Together, these data suggest that while lysosomal numbers are reduced in SMA MNs, their acidification and the activity of pH-dependent hydrolases appeared increased, particularly in the MN subpopulation that degenerates fast in culture. This raises the question of whether these abnormal acidity and protease activity in SMA MNs are due to a defective fusion of lysosomes with APs or, alternatively, whether such phenotypes are caused by a compensatory cellular mechanism attempting to overcome the dysfunctional ALP activity.

### AP-lysosome fusion is compromised in SMA MNs, however lysosomal axonal transport is not impaired

Decreased lysosomal numbers and increased lysosomal enzymatic activity are observations that, at first glance, seem counterintuitive. To shed light on this apparent conundrum, we next explored whether the overall autophagy flux is reduced in SMA MNs due to diminished lysosomal numbers, which should therefore eventually result in the formation of intracellular aggregates, or whether the increase in protease activity compensates for it and preserves proteostasis.

While we and others have previously reported an enhanced number of APs in several SMA models and human cells, (*14–18*), and interestingly, mRNA levels of *MAP1LC3B*, *UVRAG* and *GARABAP*, all key players in AP initiation, nucleation, maturation and fusion steps (*42*) appeared upregulated in both SMA types 2 and 1 neuromuscular organoids compared to their isogenic controls in our recent scRNAseq study (Figure S3J) (*19*), AP clearance has been found impaired. However, a molecular explanation for the malfunctioning AP clearance is still missing. Taking advantage of our human SMA isogenic model, we measured autophagy flux using a lentivirus (LV) carrying a mCherry-GFP-LC3 tandem reporter (LC3-TR) (*43, 44*). Using this tool, APs are visualized as both mCherry^+^ and GFP^+^ puncta, and therefore detected as yellow dots. Due to the sensitivity of GFP to acidic pH levels, the GFP signal is quenched upon AP fusion with acidic lysosomes. This allows autophagy flux to be monitored by quantifying the percentage of autophagic vesicles before (yellow APs = mCh^+^/GFP^+^ puncta) and after fusion with lysosomes (red APLs = mCh^+^/GFP^-^ puncta) (*43, 44*) (Figure 3A-B). MNs were transduced with the LC3-TR LV and the percentage of APLys was quantified 9 days after transduction. SMA type 2 MNs showed a marked decreased in the percentage of APLys (out of the total number of autophagic vesicles) when compared to its isogenic control (Figure 3A, C), and a significant reduction in APLs was observed in type 1 MNs compared to both healthy MNs (Figure 3A, D). This is the first time that an impaired autophagic flux in SMA MNs is reported in a human isogenic model, which also proves SMN causality. It is noteworthy that, due to the lentiviral approach required to label the autophagic vesicles and the time window required for the construct to be expressed, these results correspond to DIV10 MNs, and hence it is possible that a more severe phenotype is unnoticed by the loss of the most vulnerable MNs at this stage. To assess whether the reduced APLys formation was due to a deficient AP-lysosome fusion, we next combined the LC3-TR labelling with immunostaining against the lysosomal CTSB. SMA type 2 MNs showed a significant decrease in the percentage of APLs positive for CTSB compared to their isogenic control and the WT MNs, while SMA type 1 MNs showed a similar significant decrease compared to the WT and a trend for the same pattern in comparison to their isogenic control (Figure 3E-F). This indicates that, in addition to a reduced number of APLs per MN, a fusion defect between APs and lysosomes could be present in SMA MNs.

**Figure 3.**
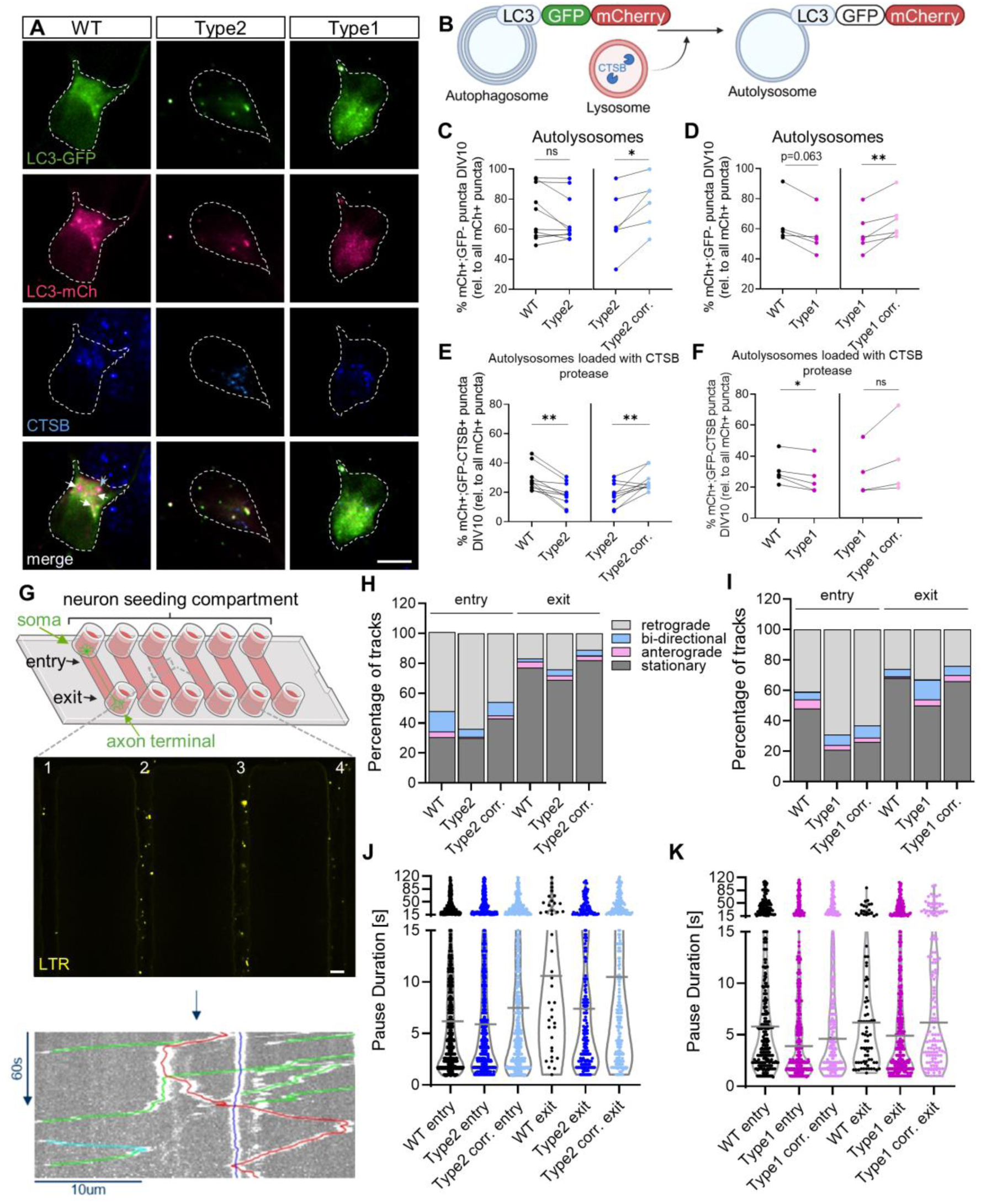
Autolysosome formation is reduced in SMA type 1 MNs, however, lysosomal axonal transport is not impaired. (**A**) Representative images of healthy and SMA MNs transduced with a lentivirus carrying a LC3 tandem reporter (LC3-TR), fixed at DIV10 and immunostaining against CTSB (blue). Scale bar, 10µm (White arrows indicate autolysosomes, blue arrowhead indicate autolysosome positive for CTSB staining). (**B**) Schematic representation of the autophagy flux measured via LC3-TR. (**C**) Quantification of the percentage of autolysosomes (mCh^+^;GFP-;LC3) out of all autophagy vesicles (mCh^+^;LC3) per neuron in SMA type 2 vs isogenic control and healthy control and (**D**) type 1 vs isogenic control and healthy DIV10 MNs. (**E**) Quantification of the percentage of autolysosomes containing the lysosomal protease CTSB (mCh^+^;GFP-;LC3;CTSB^+^) out of all autophagy vesicles (mCh^+^;LC3) per neuron in SMA type 2 vs isogenic and healthy control and (**F**) type 1 vs isogenic control and healthy DIV10 MNs (each circle represents an independent experiment, each representing the mean of n= 3 technical replicates, 2-tailed Paired t-test. Mean ± SEM is shown. Samples from the same experiment are connected by a line). (**G**) Schematic representation of a microfluidic device used to determine lysosomal trafficking along the MN axons. The magnified squared shows the entry region of 4 microchannels. Scale bar, 20µm. Bottom, example kymograph of LTR^+^ vesicle tracks segmented into runs and pauses. (**H-I**) Proportion of LTR^+^ vesicle (yellow) classified based on their movement - retrograde (light grey), bidirectional (blue), anterograde (pink) and stationary (dark grey). Number of individual lysosomes quantified are as followed: (H) WT entry n= 756, Type 2 entry n= 561, Type 2 corr. entry n= 471, WT exit n= 84, Type 2 exit n= 903, Type 2 corr. exit n= 453 and (I) WT entry n= 252, Type 1 entry n= 313, Type 1 corr. entry n= 153, WT exit n= 81, Type 1 exit n= 329, Type 1 corr. exit n= 155. Quantification of mean pause duration of each LTR^+^ vesicle in the SMA type 2 (**J**) and SMA type 1 MNs compared to isogenic and healthy controls (**K**). (Each dot represents an individual lysosome. Median is indicated).

The maturation of APs along the axon involves several vesicle fusion processes. APs fuse with increasingly acidic vesicles that contain lysosomal hydrolases and other proteins and eventually with mature lysosomes in the soma to form actively degrading APLs (*35, 43*). In addition, because neuronal APs form predominantly in the distal axon, they must undergo retrograde transport towards the cell body to fuse with lysosomes for degradation (*45, 46*). The retrograde transport of APs is mediated by the dynein-dynactin motor complex, which is recruited to APs via the Ras-related protein 7 (RAB7) along microtubules. The same complex also regulates the retrograde transport of late endosomes, lysosomes and other autophagy-related vesicles (*42*). Mitochondrial axonal transport has been reported reduced in primary SMA mouse MNs and SMA hiPSC-derived MNs (*47–50*). Similarly, the levels of several subunits of the dynein complex were found significantly altered in an RNAseq study on hiPSC-derived purified SMA type 1 MNs compared to healthy controls (*23*) as well as in SMA types 2 and 1 neuromuscular organoids compared to their isogenic controls in our recent scRNAseq study (Figure S3J) (*19*). Together, these evidence supports a dysregulation of axonal organelle transport in SMA MNs. As the direct measurement of AP-lysosome fusion is notoriously challenging, we next sought to explore whether SMA MNs were affected by a general lysosome transport defect. WT, SMA, and isogenic corrected MNs were seeded into microfluidic devices for individual LTR-labelled lysosome trafficking analysis within the motor axons (Figure 3G). These devices are composed of a somatic and an axonal compartment connected by microchannels through which the motor axons migrate. As previous studies have described that the microchannel section proximal to the soma compartment (“entry”) can contain both dendrites and axons, whereas the most distal section (900 µm distance from entry) contains exclusively axons (“exit”) (*51, 52*), lysosomal transport was analysed in both regions. The tracked lysosomes were classified, based on their movement, as retrograde, bi-directional, anterograde and stationary. Analysis revealed clear differences between SMA MNs and controls. Both SMA MN types showed a marked increase in retrogradely transported lysosomes compared to isogenic and healthy control MNs (Figure 3H-I). This was accompanied by a reduction in the percentage of stationary lysosomes, with type 1 MNs showing the most profound decrease (Figure 3H-I). This increase in the percentage of moving lysosomes correlated with the reduction in lysosomes pause duration, especially in the “exit” axonal section (Figure 3J-K). Mean velocities remained comparable across all MN genotypes (data not shown). Together, these findings indicate enhanced lysosomal trafficking from the axon terminals towards the cell bodies in SMA MNs. This suggests that the reduced AP-lysosome fusion, evidenced by decreased APLys and CTSB^+^ APLys percentages, is likely not caused by impaired lysosome transport. While this data clearly demonstrate altered lysosomal dynamics in SMA MNs compared to healthy MNs, two possible interpretations emerge: the increased transport could either represent a compensatory response to impaired degradative capacity, or it might indicate dysregulation of controlled lysosome trafficking. This distinction needs to be explored further since lysosomal pauses along the microtubules are critical for fusion between lysosomes and APs (*53*).

### Lysosomal defects upon SMN deficiency are specific to MNs and not present in cortical neurons

It is known that most neurodegenerative diseases, like AD, Parkinson’s (PD), Huntington’s (HD) and ALS, exhibit a pathology that selectively targets defined subsets of neurons. While this effect is well documented, the mechanistic origin of this selective neuronal vulnerability is still not fully understood (*54*). This phenomenon also occurs in SMA, where spinal MNs are the primary pathological target of SMN deficiency, while other neuronal types, like cortical neurons, remain mostly unaffected (*19, 55*). Furthermore, in a previous study we discovered a second level of selective vulnerability as, within the spinal MNs, the ones harbouring intrinsically low SMN protein levels died significantly faster in culture than the ones expressing higher amounts (*21*). In contrast, cortical neurons (CxNs) derived from the same SMA hiPSC lines, despite having analogous SMN protein deficiency and presenting similar cellular heterogeneity in SMN protein, did not show SMN-dependent death (*21*).

To explore whether the discovered ALP axis defects in SMA were specific to MNs and not present in other neuronal types, which could argue for an underlying cause of their selective vulnerability, we investigated several lysosomal parameters in CxNs. Our isogenic cohort of hiPSCs was subjected to a well-established protocol for CxN generation based on the inducible overexpression of *NGN2* (*56*) (Figure 4A). CxN differentiation was confirmed by immunostaining of the CxN-specific marker BRN2 (POU3F2) together with the pan-neuronal marker microtubule-associated protein 2 (MAP2). No difference in the CxN differentiation capacity between the hiPSC lines was observed (Figure 4B-C). First, longitudinal quantification of neuronal survival over two weeks in culture revealed that while MN death correlated with disease severity (SMA type 1 MNs survived worse in culture than type 2 and these worse than healthy controls), and was fully rescued in the isogenic controls (Figure S4B), SMA CxN survival was similar to that of isogenic and healthy controls (Figure S4A, C). After confirming the selective vulnerability of MNs and the resilience of CxNs to SMN deficiency, we next quantified the number of lysosomes per individual CxN using LAMP1 immunostaining and LTR labelling. Neither the lysosomal numbers nor lysosomal acidification were altered in SMA type 2 or type 1 CxNs compared to their isogenic pairs or healthy control (Figure 4D-H). Altogether, these data show that, despite carrying the same SMA-causing mutations and having the same genetic background, while MNs exhibit a SMN-dependent death and defective lysosomal biology, lysosomal numbers and acidification in CxNs, as well as their general health remained unaffected. These findings indicate that the lysosomal faults detected upon SMN protein deficiency affect specifically MNs and thus could contribute to their selective degeneration in the patients.

**Figure 4.**
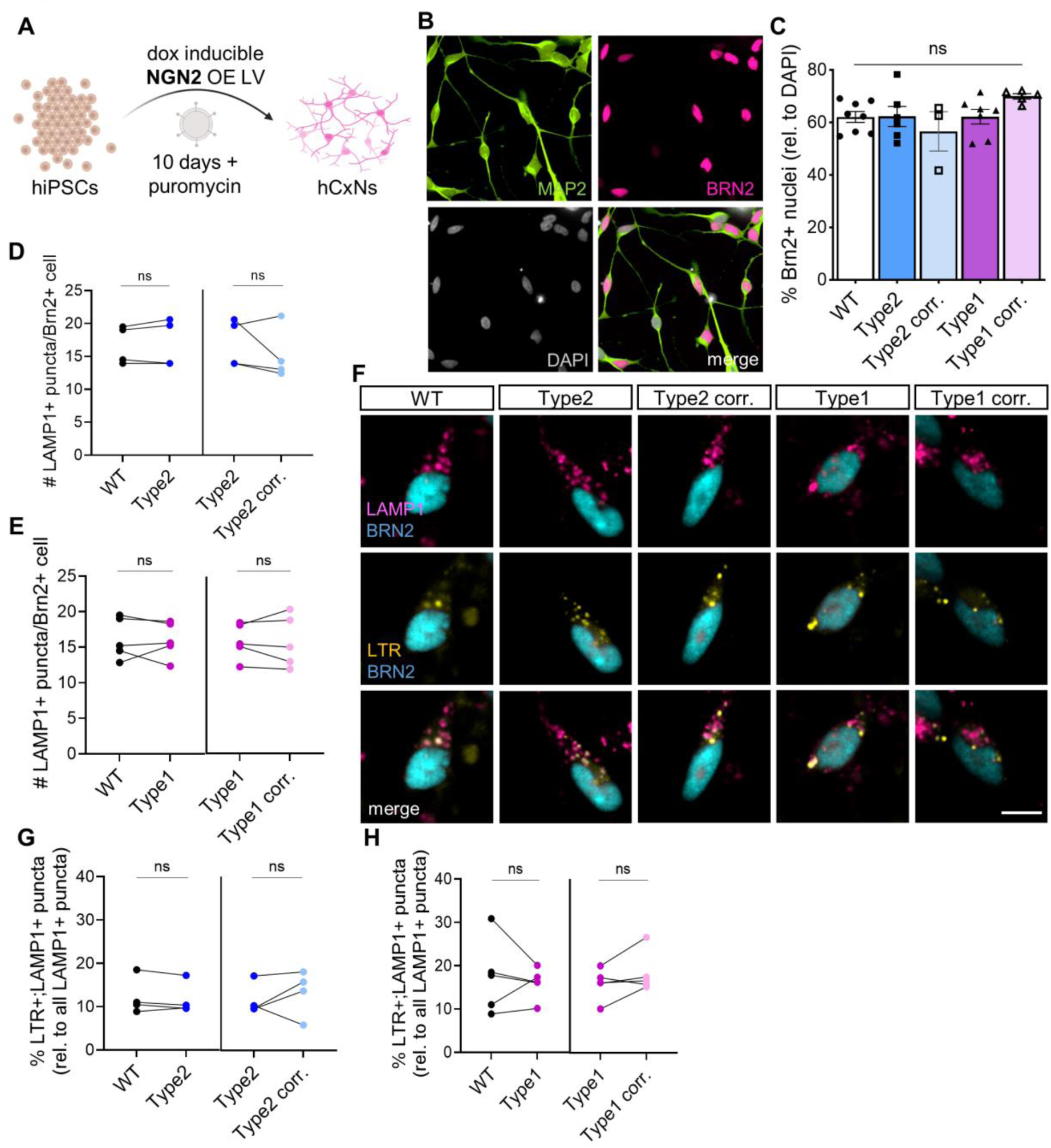
Lysosomal defects upon SMN deficiency are specific to MNs and not present in cortical neurons. (**A**) Schematic representation of the generation of cortical neuron (CxN) cultures from hiPSCs by doxycycline-inducible *NGN2* overexpression. (**B**) Representative images of CxNs 10 days after *NGN2* overexpression (Scale bar, 10µm) and quantification of the percentage of BRN2^+^ nuclei (**C**) (One-way ANOVA with Bonferroni’s multiple comparison correction. Each point represents an independent experiment, each representing the mean of n= 3 technical replicates). (**D**) Quantification of the number of LAMP1^+^ puncta per BRN2^+^ cell in SMA type 2 (**E**) and SMA type 1 MNs vs isogenic control and healthy CxNs. (**F**) Representative images of hiPSC-derived BRN2^+^ CxNs showing labeled lysosomes with LTR (yellow) and immunostaining against LAMP1 (magenta). Scale bar, 10µm. Quantification of the number of LTR^+^;LAMP1^+^ puncta per BRN2^+^ cell in SMA type 2 (**G**) and SMA type 1 CxNs vs isogenic and healthy control CxNs (**H**) (2-tailed Paired t-test. Each independent experiment is plotted as individual circle, each representing the mean of n= 3 technical replicates. Samples from the same experiment are connected by a line. Mean ± SEM is shown).

### SMN deficiency results in the formation of protein aggregates

A hallmark of many NDs is the presence and accumulation of insoluble protein aggregates that contribute to the collapse of protein homeostasis, ultimately leading to cellular toxicity (*57–59*). The class of aggregating protein is often specific to each disease, especially if this is caused by a mutation that alters the turnover, conformation, subcellular localization or distribution of the resulting protein, such as extracellular amyloid beta plaques and intracellular hyperphosphorylated tau tangles in AD, α-synuclein^+^ Lewy bodies in PD, mutated huntingtin deposits in HD, or TDP-43, SOD1 or FUS in ALS (*60*). In contrast, SMA is not caused by the accumulation of a mutated protein but instead by the deficient levels of a protein essential for MN survival, the SMN protein. Nevertheless, in our previous study we found both p62 and ubiquitinated proteins, main components of protein aggregate structures, accumulated in SMA MNs (*14*), mutations in *UBA1* cause X-linked SMA (*61*) and widespread perturbations in ubiquitin homeostasis have been also described in SMA (*62*), matching the largely increased levels of the ubiquitin gene expression (*UBC*) in SMA neuromuscular organoids found through our recent scRNAseq data set (Figure S3J) (*19*). It is well known that in NDs aberrant protein aggregation often correlates with dysfunction of the ALP (*63, 64*). Given the lysosomal and autophagy defects that we have identified in SMA MNs in this study, we explored whether SMN deficiency could result in an aberrant protein aggregation phenotype. First, as lysosomal dysfunction upon SMN downregulation was observed in HEK293T cells (Figure S1), similar cells were used to perform differential solubility assays to separate proteins into soluble and insoluble (aggregate-containing) fractions (Figure 5A). Immunoblot analysis of these fractions confirmed that SMN protein is not enriched in the insoluble fraction (Figure 5B-C), and uncovered that both the autophagy substrate and receptor and aggregation marker p62/sequestosome-1 (*65*) and the aggresome cage protein vimentin (Morrow et al. 2020) were significantly enriched in the insoluble fractions upon SMN knock-down (Figure 5B, D-E). This result aligns with previous studies showing that p62 levels are increased in SMA in vitro and in vivo models (*14, 66*). Next, these aggregation markers were measured by western blot in WT and SMA MNs. Both SMA type 2 and type 1 MNs showed increased levels of p62 and vimentin in a disease severity-dependent manner (Figure 5F-H). Aiming to validate these results using a second approach, vimentin immunostaining was performed in early and late MN cultures. Quantification of vimentin^+^ puncta per MN at DIV2 showed a significant increase in SMA compared to WT MNs (Figure 5I-K), and was not detected in the longer-surviving MNs (Figure 5L-M). Finally, as further confirmation of the presence of protein aggregates, Proteostat, an assay to detect and measure denatured proteins within aggresomes and aggresome-like inclusion bodies in fixed cells (Vasudevan et al. 2024) was utilized. The number of Proteostat^+^ puncta per MN was significantly increased in SMA MNs, specifically at DIV2 (Figure 5N-P) and not in DIV10 MNs (Figure 5Q-R). Additionally, the results showed that both the number of vimentin^+^ and Proteostat^+^ puncta generally increased in MNs of all genotypes at DIV10 compared to DIV2. This indicates that MNs accumulate undegraded protein aggregates as they survive longer periods of time, probably due to the intrinsic stressful conditions of in vitro culture, mimicking the in vivo scenario (*13*), however, a clear increase of these aggregates is only observed in SMA MNs relative to healthy controls while both vulnerable and resistant populations are still present in the culture.

**Figure 5.**
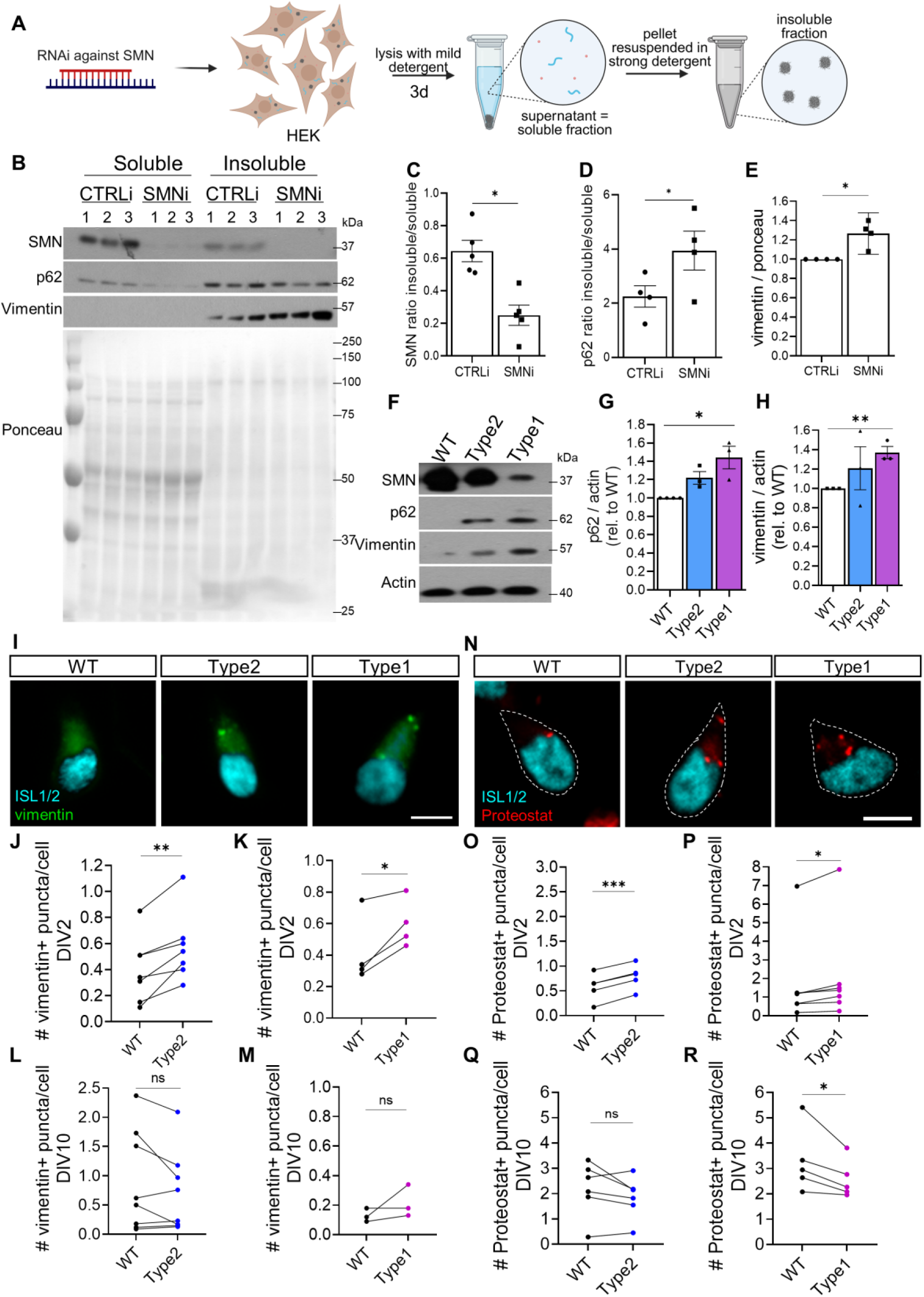
SMN deficiency results in the formation of protein aggregates. (**A**) Schematic representation of differential solubility assay to isolate soluble and insoluble protein fractions. (**B**) Representative immunoblot of soluble and insoluble protein lysates from HEK293T cells treated with scramble RNAi or RNAi against *SMN* for 72 hours. Quantification of the percentage of SMN (**C**) and p62 proteins (**D**) detected in the insoluble vs soluble lysate fractions in the control and SMN KD treated cells. (**E**) Quantification of vimentin protein levels in the insoluble fractions (N= 3-5 independent experiments, each including n= 3 technical replicates. 2-tailed Paired t-test). (**F**) Representative immunoblot from healthy and SMA hiPSC-derived MN whole protein lysates and p62 (**G**) and vimentin (**H**) protein levels quantifications (2-tailed Paired t test, N= 3-4 independent experiments). (**I**) Representative images of DIV2 SMA vs healthy MNs immunostained against ISL1/2 (cyan) and vimentin (green). Scale bar, 10µm. Quantification of the number of vimentin^+^ puncta per ISL1/2^+^ MNs in SMA type 2 vs control (**J**) and SMA type 1 vs control (**K**) MNs at DIV2 and at DIV10 (**L**-**M**, for SMA type 2 and type 1, respectively). (**N**) Representative images of DIV2 SMA vs healthy MNs immunostained against ISL1/2 (cyan) and Proteostat (red). Scale bar, 10µm. Quantification of the number of Proteostat^+^ puncta per ISL1/2^+^ SMA type 2 vs control (**O**) and SMA type1 vs control (**P**) MNs at DIV2 and at DIV10 (**Q**-**R**, for SMA type 2 and type 1, respectively) (2-tailed Paired t-test. Each circle represents an independent experiment, each representing the mean of n= 3 technical replicates. Samples from the same experiment are connected by a line). Mean ± SEM is shown.

We and others have previously reported dysregulation of mTOR activity in SMA (*14, 18, 67*). Given mTOR’s critical role in protein synthesis (*68, 69*), to rule out that the increased levels of the protein aggregation markers detected in SMA MNs were due to an abnormally increased protein synthesis, which could saturate the neurons’ proteostasis pathways and lead to protein aggregation, we measured nascent protein biosynthesis (*70*) in SMA MNs. Using this method, the results showed that overall protein synthesis is partially reduced in SMA MNs compared to healthy control (Figure S5), which is consistent with previous studies revealing a depletion of ribosomes and protein translation upon SMN loss (*71, 72*). These findings demonstrate that SMN deficiency causes a protein aggregation phenotype in vulnerable SMA MNs, which may have been previously overlooked due to the early death of these MNs.

### TFEB is dysregulated in SMN deficient cells and its overexpression ameliorates MN degeneration in vitro and in vivo

One way through which mTORC1 negatively regulates autophagy functionality is via the phosphorylation of the transcription factor TFEB, which determines its subcellular localization (*73, 74*). TFEB is the main regulator of lysosomal biogenesis, lysosomal acidification and fusion with APs, and hence it increases autophagy flux, and it is conserved through evolution (*75, 76*). mTORC1 is the main negative regulator of TFEB activity and when activated it phosphorylates TFEB at least at three serine residues (S122, S142, and S211). This promotes TFEB nuclear export or prevents it from being transported into the nucleus and retained in the cytoplasm, where it can no longer promote the expression of the CLEAR (Coordinated Lysosomal Expression and Regulation) gene network, a large cohort of lysosomal and autophagic genes crucial for the de novo formation of lysosomes and APs (*76*). Based on our results, we questioned whether the overactivated mTORC1 in SMA MNs could lead to reduced activity of TFEB and this to the observed defects in the ALP axis. To test this hypothesis, we first knocked down SMN in a HEK293T line stably expressing an inducible shRNA and measured TFEB levels in cytoplasmic and nuclear cellular fractionations. Western blot analysis revealed that SMN KD did reduce the nuclear:cytoplasmic TFEB ratio (Figure 6A-C). This data indicates that SMN deficiency impedes TFEB’s nuclear translocation, which could then lead to the observed reduction in lysosomal markers (Figures 1, 2 and 6D).

**Figure 6.**
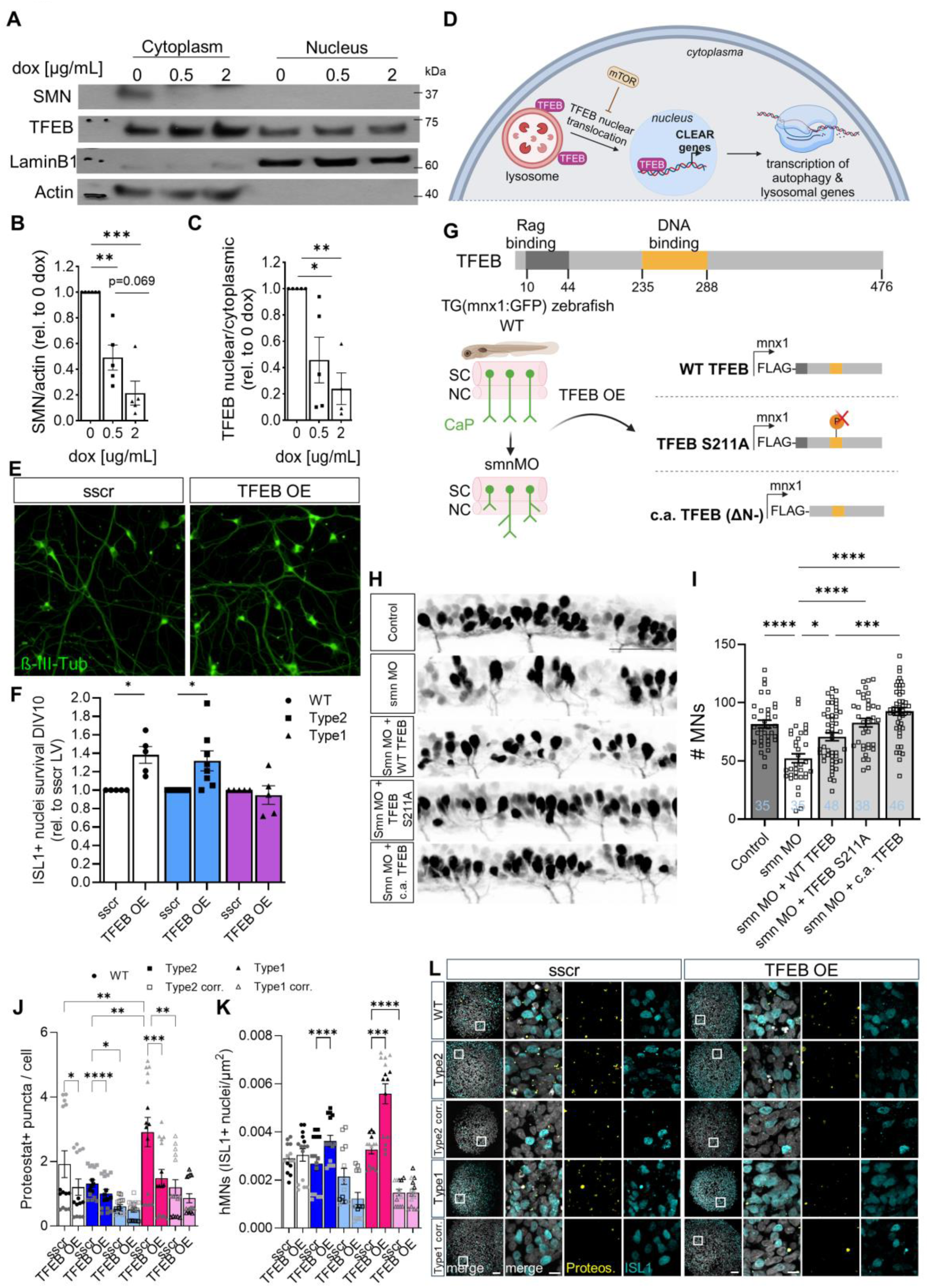
TFEB is dysregulated in SMN deficient cells, and its overexpression ameliorates MN degeneration in vitro and in vivo. (**A**) Representative immunoblot from protein lysates subjected to subcellular fractionation from HEK cells stably expressing RFP:shSMN and treated with the indicated doxycycline concentrations for 72 hours. Quantification of SMN (**B**) and (**C**) TFEB protein levels in the subcellular fractions upon SMN downregulation (One-way ANOVA with Bonferroni’s multiple comparison correction, N= 4-5 independent experiments). (**D**) Schematic representation of TFEB nuclear translocation and lysosomal functionality regulation (Biorender). (**E**) Representative images of hiPSC-derived MNs fixed and immunostaining against ß-III-tubulin (green) 10 days after being transduced with a control (sscr) or TFEB-overexpressing lentivirus (LV). Scale bar, 50µm. (**F**) Quantification of the percentage of SMA and healthy ISL1^+^ MN survival upon TFEB overexpression (2-tailed Paired t-test, each point represents an individual experiment). (**G**) Schematic representation of TFEB constructs used for over expression in a Zebrafish SMA model (Biorender). (**H**) Representative lateral views of spinal motor axons from a zebrafish SMA model 28 hours after fertilization and morpholino injection. Scale bar, 50μm. (**I**) Quantification of spinal MNs in zebrafish treated with the different conditions (Kruskal-Wallis test with Dunn’s post hoc test. n= 35 uninjected, n= 35 smn MO; n= 48 smn MO + WT TFEB-FLAG; n= 38 smn MO + TFEB S221A-FLAG; n= 46 smn MO + constitutively active (c.a.) TFEB-FLAG). Mean ± SEM is shown. (**J**) Quantification of the number of Proteostat+ puncta per cell (stained with HOECHST) on crysections from WT, SMA type 2 and type 1 and their corresponding isogenic control EBs transduced with sscr or TFEB OE LV from day 9 to day 15 of development and (**K**) quantification of the percentage of ISL1+ MNs in the same samples (N= 3 independent color-coded experiments, each containing 5 EBs. At least 2 different cryosections per EB were quantified and their average represented as an individual point. Two-way ANOVA with Tukey’s multiple comparison test). (**L**) Representative cryosections from day 15 sscr- or TFEB OE-LV transduced EBs immunostained against Proteostat (yellow) and ISL1 (cyan), nuclei stained with Hoechst (grey). Squared areas on the left panels (scale bar, 50µm) are magnified on the right-side ones (scale bar, 10µm).

It has been shown in the context of several NDs that TFEB dysregulation can act as a driver of autophagy dysfunction (*77, 78*) and that TFEB overexpression (OE) can prevent or ameliorate neurodegeneration in different model systems (*13, 79–81*). Therefore, we next examined whether TFEB could constitute a potential therapeutic target in SMA, which remained unexplored. First, mRNA levels of TFEB and multiple CLEAR genes were significantly increased in HEK293T cells transduced with a TFEB overexpressing lentivirus (TFEB LV) compared to a scramble overexpressing control (sscr LV) (Figure S6A-B). Similarly, WT MNs transduced with the TFEB LV showed an enhanced number of LAMP1^+^ puncta per MN by immunostaining (Figure S6C). This data confirmed that LV-mediated TFEB OE increased TFEB mRNA levels and expression of its target genes. Next, to test whether TFEB OE would ameliorate the observed lysosomal/autophagy pathological phenotypes in SMA MNs and enhance their survival, WT and SMA MNs were transduced with either the sscr or TFEB LVs. MN survival was determined by quantifying the number of ISL1^+^ MNs alive after 10 days in culture compared to those detected after plating (2 days in culture). Both WT and SMA type 2 MNs showed a significant increase in survival upon TFEB OE compared to the MNs transduced with the control virus (Figure 6E-F). MN axonal growth, pathfinding, and target innervation defects have been reported in different SMA models (*82, 83*). Therefore, neurite length was also measured and normalized by cell count in SMA type 2 MNs. These results revealed a significant increase in neurite length per cell upon TFEB OE specifically in the SMA MNs (Figure S6D-E). No effect on survival was detected in the severe SMA type 1 condition, potentially because these MNs undergo a significantly more pronounced cell death over the first week in culture (Figure 1B) that cannot be rescued due to the temporal window required for TFEB levels to raise after LV transduction. These data therefore shows that the positive effect of TFEB OE is not limited to diseased MNs but can also boost the survival of WT MNs. This aligns with our observation that WT MNs also display a heterogeneity of SMN levels that determines their probability of survival upon stress (*21*), and that improving ALP functionality can promote neurite outgrowth (*84*) and constitutes a pro-survival mechanism (*85, 86*).

To test whether TFEB activation would also improve SMA disease phenotypes in vivo, we overexpressed it in an SMA zebrafish model (*62*), a malleable system to determine the effects of genetic manipulations on MN health. Five different experimental conditions were tested: zebrafish embryos of the tg(mnx1:GFP) line in which MNs and axons are fluorescently labelled were (1) left uninjected as controls, (2) injected with a morpholino against smn (smnMO), (3) smnMO plus wild-type TFEB (WT TFEB), (4) smnMO plus TFEB in which serine 211 was mutated to an alanine (TFEB S211A) to prevent mTOR-mediated phosphorylation and therefore induce TFEB translocation to the nucleus (*76, 78*), and (5) smnMO plus a truncated version of TFEB that renders it constitutively active (c.a. TFEB) (Figure 6G) (*87*). All TFEB constructs were cloned under the *mnx1* MN specific promoter and were N-terminally tagged with FLAG. At 26-28 hours post-fertilization (hpf), the effect of TFEB overexpression was measured by analyzing the survival of spinal caudal primary (CaP) MNs. The quantifications showed that SMN KD, as reported before (*88*), decreased the number of MNs, while overexpression of all three TFEB constructs significantly rescued this phenotype (Figure 6H-I). In addition, immunostaining analysis demonstrated that all three TFEB constructs localized to the cell body of MNs (Figure S6F). It has been previously published that SMN downregulation in zebrafish also results in the aberrant ramification of spinal MN axons (*62, 89, 90*). An abnormally high presence of MNs with aberrant axonal branching was observed upon SMN KD compared to uninjected controls, which was significantly ameliorated in all three overexpressed TFEB conditions (Figure S6G-H). Importantly, the overexpression of a TFEB construct lacking the DNA-binding domain did not ameliorate the pathological MN axonal branching in the SMA zebrafish, which indicates that TFEB transcriptional activity is essential for its cytoprotective function (Figure S6I-J). Together, these data indicate that the induction of lysosome/autophagy functionality through TFEB overexpression not only increases the survival of MNs in our human *in vitro* model but also improves MN health *in vivo* in an SMA Zebrafish model.

As our data showed that SMN MNs undergo the accumulation of undegraded protein aggregates, which are widely linked to neuronal death in virtually all neurodegenerative diseases (*12, 91*), likely caused by the impaired APLys functionality, we next investigated whether TFEB overexpression could also ameliorate this common neurodegeneration hallmark. To increase the window of time of enhanced TFEB levels and thus the probability of it exerting its cytoprotective role before the most vulnerable MNs were lost, next we overexpressed TFEB on day 9 EBs, when MNs start to differentiate (*19*), and fixed them on day 15. High-content imaging and automated quantification of Proteostat^+^ protein aggregates in EB cryosections showed that SMA type 1 EBs had a greater amount than their isogenic control and than SMA type 2 and healthy controls (Figure 6J, L), confirming our results on dissociated MNs (Figure 7N-R). In addition, TFEB OE significantly reduced the number of Proteostat^+^ puncta compared to the EBs transduced with the sscr-control LV, especially in SMA type 1 (Figure 6J, L). This reduction in the abundance of protein aggregates mediated by TFEB OE was accompanied by a notable increase in the number of ISL1+ MNs in the SMA type 1 and type 2 EBs (Figure 6K-L).

**Figure 7.**
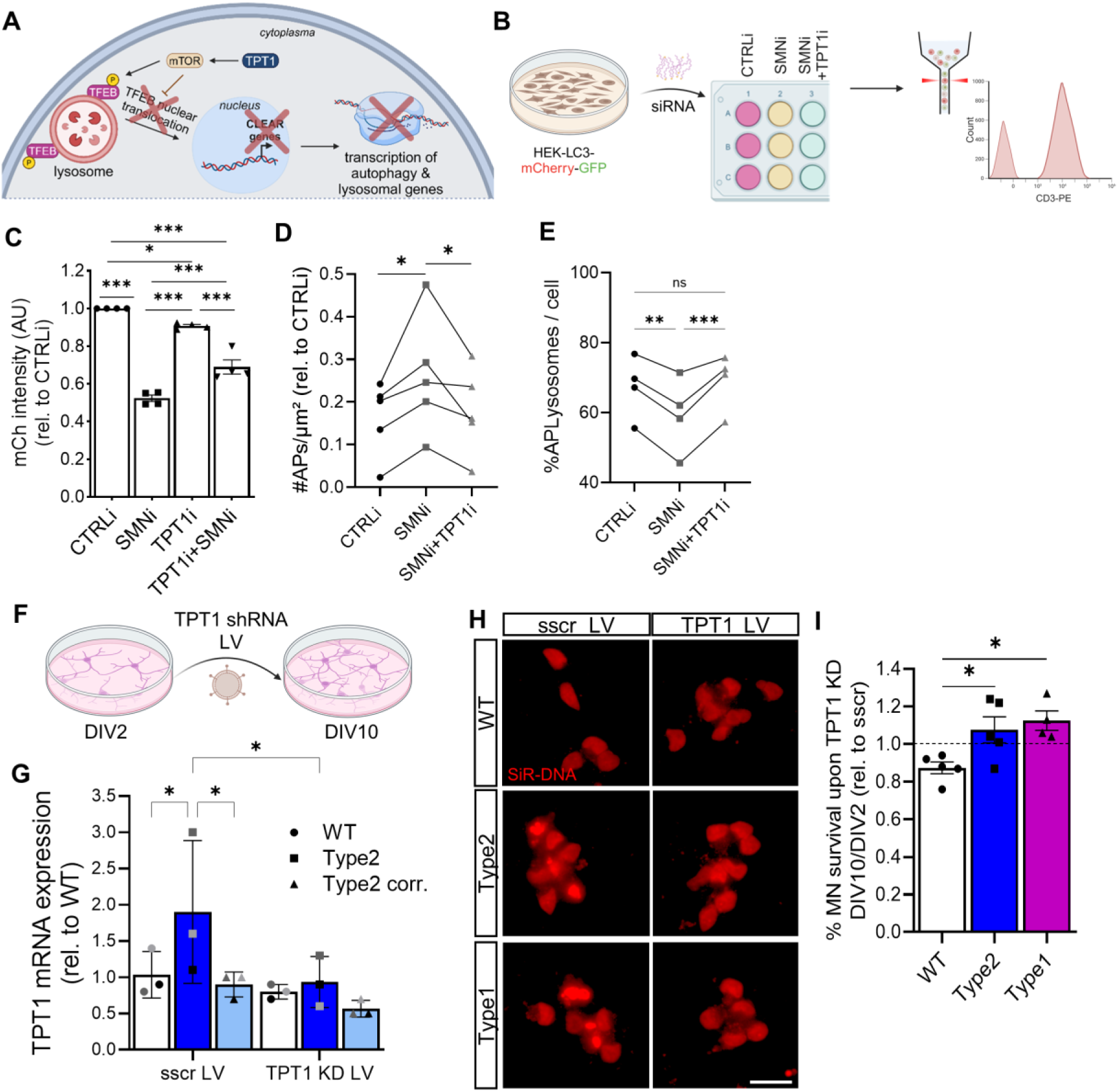
TPT1, a potential link between SMN reduction, mTOR activation, TFEB dysregulation and lysosomal/autophagy defects in SMA. (**A**) Scheme indicating the ALP functionality impairment potentially caused by an increased activity of TPT1 in SMA MNs (Biorender). (**B**) Scheme indicating the experimental design followed in C. (**C**) Quantification of mCherry fluorescence intensity by FACS analysis of a stably transduced HEK-LC3-TR line subjected to RNAi for 3 days to knock-down of the indicated targets (One-way ANOVA with Bonferroni’s multiple comparison correction, N= 4 individual experiments, each representing the mean of n= 3 technical replicates). (**D)** Quantification of the number of APs (mch^+^;GFP^+^) per µm² and (**E**) the percentage of autolysosomes (mCh^+^;GFP-) per cell stably transduced HEK- LC3-TR cells subjected to *SMN* or *SMN*+*TPT1* knockdown (2-tailed Paired t-test, each circle represents an independent experiment, each representing the mean of n= 3 technical replicates. Samples from the same experiment are connected by a line). (**F**) Scheme showing lentiviral TPT1 knockdown in MNs. (**G**) Quantification of *TPT1* mRNA expression levels measured by qPCR from WT and SMA types 2 and 1 MNs after *TPT1* knock-down (N= 3, each representing 2 technical replicates. Two-way ANOVA with Fischer’s LSD test). (**H**) Representative images of DIV10 SMA vs healthy MNs treated for 9 days with sscr or *TPT1* shRNA carrying LVs labelled with the live dye SiR-DNA (red). Scale bar, 20µm. (**I**) Fold change survival of hiPSC- derived ISL1^+^ MNs treated for 9 days with *TPT1* shRNA LV relative to the survival of sscr- transduced MNs (One-way ANOVA, N= 5 individual experiments, each representing the mean of n= 3 technical replicates). Mean ± SEM is shown.

Lastly, we explored whether a combinatorial treatment approach composed of a SMN-increasing compound together with TFEB OE, would lead to an improved outcome on SMA MN survival. We thus treated SMA type 2 MNs simultaneously with risdiplam, one of the three approved treatments for SMA patients (*1, 92*) and our TFEB OE LV for 9 days (Figure S7A-B). Unexpectedly, while risdiplam (75-600 nM) alone did increase average SMN levels by 30-40% compared to control-treated SMA type 2 MNs (Figure S7C), it reduced their survival by 20% in all the concentrations tested (Figure S7D). The same toxic effect was observed in MNs treated with risdiplam and TFEB OE simultaneously; however, even under these unfavourable conditions, TFEB OE prevented MN death compared to the sscr LV-treated MNs (Figure S7E). Taken together, our data indicate that dysregulation of TFEB localization triggered by deficient SMN protein levels could constitute the underlying cause for the impaired ALP in SMA MNs. Our results thus also suggest that increasing TFEB levels has the potential to slow down or even prevent SMA MN degeneration and loss.

### TPT1 upregulation as a potential link between reduced SMN levels and mTORC1 activation in SMA

We next decided to search for a mechanistic link between SMN protein deficiency and mTOR overactivation, causing the TFEB dysregulation and ALP failure that we have discovered in SMA MNs. A promising candidate was identified from our RNAseq on FACS-purified SMA type 1 vs healthy control MNs (*23*), *TPT1* (Tumour Protein, Translationally-Controlled 1). *TPT1* is a ubiquitously expressed gene which encoded protein regulates cellular growth, protein synthesis, cell proliferation and apoptosis (*93, 94*). Although little is known about the functionality of this protein, a study identified a role for TPT1 in the regulation of autophagy by activating mTOR, as the reduction of TPT1 levels led to the induction of autophagy flux via mTOR inactivation (*94*). TPT1 appeared significantly increased in SMA type 1 MNs compared to healthy controls in our RNAseq dataset (*23*) and, importantly, also significantly upregulated in the MN clusters from our recent scRNAseq on SMA type 2 and type 1 neuromuscular organoids compared to their isogenic controls (Figure S3J). We thus hypothesized that an increased TPT1 expression could be at least partially responsible for the mTOR overactivation observed in SMA MNs (*14*), resulting in the cytoplasmic sequestration and inactivation of TFEB that led to the impaired lysosomal function (Figure 7A). Should TPT1 constitute the missing mechanistic link between pathologically low SMN protein levels and ALP malfunctioning, *TPT1* knockdown should restore the defective autophagy flux and increase SMA MN survival. To test this possibility, we reduced either *SMN, TPT1* or both simultaneously via an RNAi approach on a stable HEK293T cell line expressing the LC3-TR. Three days after RNAi transfection, cells were analyzed by FACS to measure APLs mCherry fluorescence intensity as a proxy for autophagy flux efficiency (Figure 7B). mCherry signal was reduced to 50% in SMN KD cells, matching our automated image-based APLs quantification in SMA type 1 MNs (Figure 3D-F), while TPT1 KD alone only reduced mCherry fluorescence by 10% (Figure 7C). Importantly, simultaneous knockdown of both SMN and TPT1 showed a significant 20% increase in APLys mCherry signal compared to SMN KD alone (Figure 7C). This indicates that lowering TPT1 levels partially ameliorates the reduction in APLs formation caused by SMN deficiency. To confirm these results, automated imaging and quantification of autophagic vesicles on similar cell conditions was conducted. The analysis validated an increase in the number of APs and a reduction in the percentage of APLs upon SMN KD, indicative of defective autophagy flux, both of which were rescued to control levels upon additional TPT1 downregulation (Figure 7D-E). TPT1 KD did not alter SMN mRNA levels (Figure S8A) and, in addition, while SMN KD cells showed reduced *LAMP1*, *CTSB* and *TFEB* gene expression levels, TPT1 downregulation drastically corrected those changes back to control levels (Figure S8B-E).

Finally, we sought to determine whether TPT1 KD would have a similar positive effect on SMA MNs. SMA type 2 MNs showed increased phosphorylated levels of the mTORC1 downstream targets S6K1 and S6 when compared to their isogenic control and healthy control MNs (Figure S8F-I) and SMA MNs transduced with a LV carrying a shRNA against TPT1 displayed notably reduced levels of p-S6K1 protein compared to sscr-LV treated cultures (Figure S8G-H). These data confirm the reported mTORC1 pathway activation in SMA MNs (*14, 18, 67*) and indicates that TPT1 is a positive mTORC1 regulator in MNs. *TPT1* mRNA quantification from SMA type 2 MNs compared to their isogenic and healthy controls confirmed the higher expression in the diseased cells (Figure 7G) and a significant decrease in *TPT1* mRNA levels upon LV-mediated knockdown. We then quantified survival of the TPT1 KD MNs over the ones exposed to the sscr LV in healthy, SMA type 2 and type 1 cultures. Although the TPT1 shRNA-LV used did not contain any reporter or selection cassette to enable determining the percentage of neurons transduced with the construct, and in addition, a large fraction of the most vulnerable MNs could die before TPT1 expression is reduced, live imaging and automated quantification showed that while WT TPT1 KD MNs displayed a minor decrease in survival compared to the sscr-treated ones, both SMA MN types showed a mild yet significant increase upon TPT1 KD (Figure 7 H-I). Overall, these data shows that an increase in TPT1 levels induced by the loss of SMN constitutes a highly interesting target for further studies, as TPT1 downregulation ameliorates ALP alterations and enhances SMA MN survival.

## Discussion

This study provides empirical support for the hypothesis that SMN protein regulates the functionality of the autophagy-lysosome pathway (ALP). SMN deficiency impairs the nuclear translocation of TFEB, the master regulator of lysosomal biogenesis and autophagy. As a result, neurotoxic protein aggregates build up, contributing to MN degeneration in SMA. Using a hiPSC isogenic model that we recently generated (*19*) composed of one SMA type 2, one SMA type 1 patient-derived iPSC line and their corresponding isogenic corrected control, and an additional healthy control hiPSC line to generate spinal MNs, we show that SMA MNs contain a reduced number of lysosomes and autolysosomes compared to their isogenic and healthy controls. Surprisingly, the remaining lysosomes displayed increased acidity and hydrolase activity, likely as a compensatory mechanism to cope with defective AP clearance. These phenotypes are accompanied by protein aggregates, hallmark of late-onset NDs. Interestingly, these features are only present in early MN cultures that we demonstrate contain low-SMN expressing MNs, prone to die a few days after being plated (“vulnerable”), and not in high-SMN expressing ones that survive for significantly longer periods of time (“resistant”) and phenotypically resemble the isogenic and healthy control MNs. Furthermore, these disease phenotypes were absent in cortical neurons generated from the same disease hiPSCs, suggesting that ALP dysfunction contributes specifically to MN degeneration in SMA. To explore therapeutic avenues complementary to current SMN-increasing treatments, we found that lentiviral-mediated TFEB overexpression in 2D human MN cultures or spinal cord spheres significantly reduced protein aggregates and rescued MN death, especially in the severe SMA type 1 condition. Importantly, this cytoprotective effect was replicated in a zebrafish SMA model. There, we demonstrated that preventing TFEB’s DNA-binding capability abolished its rescue effect on MN axonal defects and MN survival, reinforcing its role in mediating CLEAR gene expression and lysosomal biogenesis. Furthermore, we propose the upregulation of TPT1 in SMA MNs, a positive mTORC1 regulator primarily linked to cancer, as a key link between deficient SMN protein levels and mTORC1 overactivation, resulting in reduced TFEB activity, insufficient lysosomal biogenesis and faulty ALP functionality contributing to MN death in SMA.

SMA differs from classic proteinopathies, such as AD, HD and PD, which are characterized by neurotoxic aggregates like Aβ, Tau, or mutant huntingtin in that SMA’s pathophysiology is caused by the lack of a protein, SMN, rather than by toxic protein accumulation. Additionally, as an early-onset disease, it was expected that SMA’s vulnerable MNs would succumb long before accumulating undegraded material. Consequently, proteostasis, specially the ALP, has been largely overlooked in SMA research. However, studies using mouse ESC-derived, human iPSC-derived, and SMA mouse MNs have revealed impaired AP clearance due to defective autophagy flux (*14–16*), a common ND feature. In addition, we and others reported hyperactivated mTORC1 signaling, increased p62 levels -which selectively targets SMN protein for autophagic degradation- (*14*), and direct involvement of SMN in stress granule degradation via autophagy, in coordination with C9ORF72 and p62 (*95*). These findings highlight the complex interplay between SMN protein and ALP functionality.

Lysosomes are central hubs for degrading autophagic and endocytic components, preserving neuronal survival and function. While many NDs exhibit defects in early ALP stages -such as AP formation and cargo recognition in ALS/FTD, cargo capture and degradation in PD, and AP maturation in AD-defects in later stages, including AP-lysosome fusion and lysosome acidification, are primarily linked to PD and AD (*91, 96, 97*). In this study, we have discovered that in SMA, despite having completely different early-onset etiology and clinical manifestations to PD or AD, MNs displayed impaired AP clearance due to defective lysosomal biogenesis, mediated by a dysfunctional TPT1-mTOR-TFEB axis. This unexpected similarity suggests that SMA may share systemic characteristics with late-onset NDs. Such finding is supported by recent evidence where proteomic analyses of SMA mouse spinal cords identified APP, PSEN1, HTT, MTOR, SOD1 or TDP-43 as upstream regulators of dysregulated proteins (*98*), all key players in NDs.

TFEB dysregulation has been proposed as a driver of ALP dysfunction across several NDs, including PD, AD, HD and SBMA (*73, 79, 99–103*). Under basal conditions, TFEB resides in the cytoplasm and translocates to the nucleus under stress (*73*). SMA’s lysosomal stress mechanism, neglected until now, may involve neuron-specific post-translational modifications (PTMs) and protein-protein interactions that hinder TFEB’s ALP activation. Phosphorylation by mTOR is a key regulatory mechanism keeping TFEB cytoplasmic, but other kinases, such as ERK2, AKT and GSK3β, are also involved. Consistent with this, we and others have reported that SMA MNs exhibit mTORC1 overactivation (*14, 18*) and dysregulated ERK2, AKT and GSK3β activity (*104–106*). Intriguingly, mTORC1’s activity in SMA appears tissue-specific - low in skeletal muscle but high in MNs-mirroring observations in other NDs like SBMA (*100, 107, 108*). Although the three approved SMN-increasing therapies (nusinersen, onasemnogene abeparvovec, risdiplam) have transformed SMA prognosis, these remain lifelong treatments instead of cures, with variable efficacy and potential severe adverse effects (*92, 109, 110*). Additionally, prenatal treatment might be required (*111*). This underscores the need for ongoing research to develop safer, more universally effective combinatorial treatments (*112*). Numerous studies have reported that TFEB overexpression or activation prevents the selective loss of neuronal populations in NDs by enhancing the autophagic clearance of toxic protein aggregates and improving the ALP functionality (*73, 113*). It is worth noting, though, that autophagy activation is not universally neuroprotective. In diseases with lysosomal dysfunction, inducing AP formation may worsen phenotypes by exacerbating intracellular traffic jams (*97, 102*). In SMA, it was proposed that autophagy causes MN death (*114*). In that study, autophagy inhibition via 3-MA in SMA mouse neonates ameliorated disease phenotypes. These results, however, align with our findings on an impaired ALP functionality, in which case slowing down AP formation might indeed have some beneficial effect. Clinical trials exploring TFEB activation in NDs (via AAV vectors, trehalose, or metformin) (*115, 116*) and our findings on human MNs and a zebrafish SMA model further support TFEB activation as an emerging therapeutic target to prevent MN axon degeneration and MN death in SMA.

Although the origin of the selective MN death that characterizes SMA is still a matter of intense investigation, it likely arises from splicing defects in mRNAs critical for MN survival and neuromuscular junction maintenance due to SMN loss. For example, Pellizzoni’s team demonstrated that SMN deficiency disrupts *MDM2/MDM4* splicing, critical repressors of p53, leading to p53-mediated MN death (*117–119*). Non-splicing functions of SMN, including actin binding (associated to neurite outgrowth, axonal pathfinding and synapse formation) (*120*) and ribosome-associated transcription modulation (*71*) have also been proposed. The lysosomal defects discovered in this study seem to be rooted in the abnormally increased activity of the regulator of cellular growth and autophagy inhibitor TPT1. *TPT1* mRNA alternative splicing results in multiple transcript variants encoding different isoforms (*121*) for which nothing is yet known. It is therefore possible that the defective lysosomes and impaired ALP function in SMA originate from *TPT1* mis-splicing and isoform dysregulation, though we cannot rule out SMN’s protein direct role in lysosomal functionality.

While our study utilized diverse lysosome detection approaches, such as LAMP1, CTSB, LTR, and MagicRed probes, these label acidic organelles beyond lysosomes, such as late endosomes (*122*). As defects in the endocytic pathways/endosomal trafficking have been reported in SMA MNs (*20, 123, 124*), a systematic analysis of lysosomal and endosomal markers (for degradative and non-degradative organelles, respectively) is warranted to pinpoint where the pathology first manifests and to examine potential alterations in specific cohorts of these organelles to elucidate the mechanistic link between the impaired ALP and endosomal systems.

After decades of autophagy research, its centrality to brain health and ND pathogenesis is clear. Yet, its role in SMA remains understudied. Further investigation is needed to determine the reliance of MNs on ALP, disease-stage-specific autophagy dysfunction, and the pathological cascade following ALP disruption. Despite these yet open questions, our findings suggest that enhancing lysosomal and ALP function through TFEB activation or overexpression could restore cellular homeostasis and slow SMA progression.

## Materials and Methods

### Study approval

All experiments involving hiPSCs were performed in accordance with the ethical standards of the institutional and/or national research committees, as well as the 1964 Helsinki Declaration and its later amendments and approved by the Ethics Commission at the Technische Universität Dresden. The healthy BJ siPSC (HVRDi005-A code from the European Human Pluripotent Stem Cell Registry and the SMA (HVRDi015-A, HVRDi017-A, HVRDi016-A) hiPSCs were kindly provided by Lee L. Rubin (Harvard University). Isogenic corrected lines were previously generated in our laboratory (*19*).

### Human spinal MN differentiation

hiPSCs were grown on Matrigel (Corning, 354234) coated cell culture treated dishes, maintained with mTeSR1 (STEMCELL Technologies, 85850) and split using ReLeSR (STEMCELL Technologies, 100–0484). Initiation of the MN differentiation protocol was started by dissociation of iPSCs with ReLeSR and subsequent culture in mTeSR as EBs for 3 days in ultra-low attachment dishes. For the first 24 hours mTeSR was supplemented with 10µM Y-27632 (Hölzel, M1817-10mg) and 10ng/mL FGF-2 (Millipore, 32160702). On day 4 the media was changed to neuronal induction media (NIM), containing 50% Advanced DMEM/F12 (Thermo Fisher Scientific, 12634028), 50% Neurobasal media (Thermo Fisher Scientific, 21103049), 1% Pen/Strep (LifeTechnologies, 15140122), 1% GlutaMAX (Thermo Fisher Scientific, 35050087), 0.1mM 2-ß-mercaptoethanol (LifeTechnologies, 21985023), 0.5X B27 (Thermo Fisher Scientific, 17504044), 0.5X N2 (Thermo Fisher Scientific, 17502048), 0.16% D-glucose (Carl Roth, X997.1) and 20µM ascorbic acid (Sigma Aldrich, A4403). From day 4 to day 7 NIM media was supplemented with 10µM SB 431542 (BioTechne, 1614/10) and 100nM LDN 193189 (Hölzel, M1873) to induce neural differentiation. From day 6 to day 19 1µM retinoic acid (Sigma Aldrich, R2625) and 1µM smoothened agonist (Millipore, 566660) were added. From day 11 5ng/mL BDNF (R&D, 248-BD-025/CF), from day 13 10µM DAPT (Tocris, 2634/10) and from day 15 5ng/mL GDNF (R&D, 212GD0/50CF) and 2µM cytosine arabinoside (Sigma Aldrich, R2625) were added. On day 19 the EBs were dissociated with papain/DNase solution (Worthington, LK003178, LK003172) and 8*104 cells were plated in each well of 50µg/mL poly-D-lysine (Sigma Aldrich, A-003-E), 3µg/mL laminin (Thermo Fisher Scientific, 23017015) coated p96 half-well plates (Corning, 4680). For 63X imaging cells were seeded onto glass coverslips following the same protocol. Media for culturing dissociated cultures was Neurobasal containing 1% Pen/Strep, 1% GlutaMAX, 1X non-essential amino acids (Life Technologies, 11140050), 0.5X B27, 0.5X N2, 0.16% D-glucose, 20µM ascorbic acid, 25µM 2-ß-mercaptoethanol, 2µM cytosine arabinoside, 10ng/mL BDNF and 10ng/mL GDNF. A full media change was performed 2 days after plating, then half media changes were performed every second to third day.

### Human cortical neuron differentiation

CxNs were obtained by 2-dimensional (2D) cell differentiation through lentiviral overexpression of neurogenin-2 (*NGN2*), as published before (*21, 56*). Detachment of iPSCs was performed using accutase (Corning, 25-058-Cl) to obtain a single-cell suspension. Obtained suspensions were then centrifuged at 1000rpm for 5min and the cell pellet resuspendend in CxN media consisting of Neurobasal medium with 1% Pen/Strep, 1% GlutaMAX, 1X non-essential amino acids, 0.5X N2, 0.5X B27-vitamin A (Thermo Fisher Scientific, 12587010), 20 µM ascorbic acid and 0.1 mM 2-ß-mercaptoethanol, 10 ng/ml BDNF and 10 ng/ml GDNF. After resuspension cells were counted and seeded into Matrigel coated p96 well plates (Corning, 655090) with a cell density of 4000 cells per well in mTeSR. The following day, the rtTA-containing lentivirus and NGN2-overexpressing lentivirus (*56*) were added at 1:100 each. Doxycycline treatment (0.2 µg/ml, MedChemExpress, HY-N0565B/CS2890) was started with the infection of the cells on DIV1 and refreshed every day for 3 days, then every second day until cells were fixed. From day 3 0.5µg/mL puromycin (Gibco, A11138) and 0.2mg/mL laminin were added and refreshed every 2 days. Starting at day 6 2µM cytosine arabinoside was added, and a half media change performed every 2 days.

### Human MN and CxN survival assay using sirDNA

Survival of the culture was assessed by plating cells in 96 well plates at a density of 8*104 cells/well. 3 wells per cell line were stained using the nuclear marker dye SiR-DNA (Tebu, SC007) at 125nM for 1.5h on day 2 after dissociation (MNs) or 6 days after lentivirus transfection (CxNs). Staining was repeated every 3 days with 62.5nM SiR-DNA dye for 1h. After the staining, media in the wells was replaced by fresh media. To follow survival of the neuronal cultures, stained wells were imaged every 2 days using the automated Operetta CLS microscope (PerkinElmer) at 40X magnification. Image quantification was performed using the Columbus Analysis System (PerkinElmer) to identify living cells via dye intensity and nuclear morphology.

## Cell lines

HEK293T, HEK-RFP-shSMN and HEK-LC3-TR were cultured in DMEM (Thermo Fisher Scientific, 41966052) containing 10% heat-inactivated FBS (Thermo Fisher Scientific, A5256701) and 1% Pen/Strep and were grown at 37°C with 5% CO2. For maintenance HEK cells were grown in uncoated 10cm cell culture dishes and split upon reaching ∼90% confluency. For splitting cells were washed once with PBS (Thermo Fisher Scientific, 14190169) and incubated with TrypleE (Gibco, 12605-010) for 2-3 min until all cells detached with gentle tapping. The reaction was subsequently stopped by adding the 3-fold volume of culture medium compared to the TrypleE volume. Single cell suspensions were seeded into a fresh dish with a 1:5 or 1:10 dilution. For experiments HEK cells were seeded into p24 well plates, coated with 0.1% gelatine (Millipore, ES-006-B) for 10min. Cells were detached in the same way as described above but for experiment seeding, cells were collected in a 15mL Falcon tube, centrifuged at 1000rpm for 5min, supernatant discarded and the cell pellet resuspended in 3mL of growth medium. Cell density was calculated and cells seeded with a density of 0.1*10^6^ cells per well. SMN KD in HEK-RFP-shSMN cells was induced by addition of 0.5, 1 or 2µg/mL doxycycline to the growth media starting the day after seeding. After 3-4 days cell were collected for western blot analysis.

### Cell culture treatments

For SMN KD approaches with the HEK-TRIPZ-shSMN line HEK cells were treated with the indicated concentration of doxycycline, diluted in diH2O for 72-96 hours. Doxycycline treatment was refreshed every 2 days. For live imaging procedures cells were stained with either Magic Red (BioRad, ICT937), lysosensor (Thermo Fisher Scientific, L7535), CalceinRed (Invitrogen, C34851) or LTR (Invitrogen, L7528) for 30 min in growth medium of the original solution for 30min. Cells were washed once with medium before imaging under live imaging settings and subsequent fixation. Risdiplam (Biomol, Cay29028-5) was diluted in chloroform to a concentration of 2.5mM. Cells were treated with indicated concentrations (dilutions in growth medium) for 9 days. Risdiplam treatment was refreshed every 2 days. To stain MNs with the Proteostat kit (ENZO, ENZ-51035-0025) the manufacturer’s protocol was followed. Briefly, MNs were grown in half well p96-well plates and fixed as described below. 50µL permabilization solution (0.5% triton X-100 (Sigma Aldrich, X-100), 3mM ETDA (Thermo Fisher Scientific, 15575020, pH 8.0) were added per well/plate and placed on ice with orbital agitation for 30min. After washing wells twice with PBS, normal staining procedures were followed. After removing the secondary antibody and washing with PBS the Proteostat Detection reagent was prepared by mixing 1mL of 1X assay buffer with 1µL of the Proteostat Dye. Cells were incubated for 30min at room temperature (RT) and subsequently washed 3times for 10 minutes in 1x PBS before imaging.

### Transfection and lentiviral transduction

*RNAi approach in HEK293T.* For western blot and qPCR SMN KD experiments, HEK293T cells were treated with RNAiMax lipofectamine (Thermo Fisher Scientific, 13778150) in Opti-MEM (Life Technologies, 31985062) using either a CTRL RNAi (QIAGEN, 1027280) or SMN RNAi (Santa Cruz Biotechnology, sc-36510) one day after seeding. Each p24 well of HEK293T cells containing 500µL of growth media received a mix of 50µL Opti-MEM, 3µL RNAiMAX lipofectamine and 1µL of RNAi.

On the next day 500µL of growth media were added to all wells. 3-4 days after transfection HEK293T cells were collected for western blot analysis or qPCR. For FACS analysis and qPCR experiments using the HEK-LC3-TR line, cells were treated with the RNAiMax lipofectamin protocol as described above with the CTRL RNAi, SMN RNAi, TPT1 RNAi (Santa Cruz Biotechnology, sc-40675) or a combination of SMN and TPT1 RNAi. *LC3-TR lentiviral transduction of human MNs.* The day after dissociation MNs were incubated with the mCherry-GFP-LC3 construct (LC3-TR) using a dilution of 1:100 in 50µL of media. The lentiviral construct was kindly provided by Ana M. Cuervo (Albert Einstein College of Medicine, New York, USA) and previously published (*14, 43*). The following day a complete media change was performed before returning to the standard media changes every 2-3days. *sscr and TFEB lentiviral transduction of human MNs and HEK293T cells.* The day after dissociation MNs were incubated with either lentivirus carrying a scramble sequence (sscr LV; custom made VectorBuilder vector) or the TFEB lentivirus expressing TFEB (TFEB LV; custom made VectorBuilder vector). The following day a complete media change was performed before returning to the standard media changes every 2-3days until DIV10. *TPT1 KD in human MNs.* The day after dissociation MNs were incubated with either the CTRL lentivirus (sscr LV; custom made VectorBuilder) or the TPT1 KD lentivirus (TPT1 LV; shRNAhTPT1-TRCN0000147481, Sigma). The following day a complete media change was performed before returning to the standard media changes every 2-3days until DIV10.

### RNA-isolation, cDNA-synthesis and RT-qPCR

hiPSCs, HEK cells or MNs were washed 2x PBS. Total RNA was extracted with the TRIzol reagent (Life Technologies, 15596026), and the concentration was measured with the NanoDrop TM 1000 Spectrophotometer (Thermo Scientific). 0.5–1 μg RNA was subjected DNase I digestion (Thermo Fisher Scientific Scientific, EN0521) according to the manufacturer’s instructions. RNA was reverse transcribed with High-Capacity cDNA Reverse Transcription Kit (Applied Biosystems, 4368814). RT-qPCR was performed with GoTaq qPCR Master Mix (Promega, A6002) and a Quantstudio5 Real-Time PCR Detection System (Thermo Fisher Scientific). mRNA expression levels were normalized to the expression of the human housekeeping gene h18S and relative values were determined with the comparative ddCT method. Ultimately, the gene expression was normalized against the control group of each experiment. Primers used are depicted in Supplemental Table 1.

### Immunocytochemistry and image analysis

Cells were fixed at room temperature in 4% PFA in PBS for 12-15 minutes. After fixation, cells were washed 3 times with 1X PBS. Next, permeabilization and blocking of cells was carried out by incubating and orbitally shaking cells for 20min with 0.25% TritonX-100 and 1 % Normal Goat Serum (Thermo Fisher Scientific, 31872) in 1X PBS (Perm/Block buffer). Primary antibodies were diluted in Perm/Block buffer and incubated at 4°C overnight with orbital shaking. On the next day, cells were washed 3x with 1X PBS for 10minutes with orbital shaking. Next, secondary antibodies were diluted in Perm/Block buffer and incubated for 1 hour at RT and subsequently cells were washed 3x with 1X PBS for 10 minutes with orbital shaking. In the second wash step Hoechst (Invitrogen, H3570) was added for nuclear staining, and cells were stored in 1X PBS before imaging in the Perkin Elmer Operetta CLS with the 40X water objective (25 fields per well for live imaging and at least 40 fields for fixed cells) or the Zeiss Spinning Disk microscope with AxioCam 506 mono, Monochrome CCD camera, detector element size 4.54µm, objective 63X/1.4 Plan-Apochromat, Oil. Live imaging procedures were carried out in the Operetta CLS system utilizing the heating and CO2 function (37°C, 5% CO2). Image analysis was carried out using the automated Columbus software (PerkinElmer).

### Western Blot

HEK cells or MNs were washed once with PBS and then lysed in RIPA (Thermo Fisher Scientific, 89900) buffer, containing 2% of SDS (Fisher Scientific, BP2436) and 1X HALT protease inhibitor cocktail (Thermo Fisher Scientific, 1861278). Lysed samples were incubated on ice for 20min and subsequently boiled at 95°C for 10 min before BCA measurement. *Protein concentration measurement.* Protein concentrations of cell lysates were measured using the BCA protein kit (Thermo Fisher Scientific, 23227) following the manufacturers protocol. A BCA standard curve was prepared in deionized, distilled H20 (diH2O) ranging from 0.06 to 2mg/mL. Duplicates of each standard and sample were added to a clear p96 well plate and 190µL of the reaction mix added. The BCA reaction mix was prepared by mixing solution A and solution B in a 50:1 ratio. Plates were then incubated at 37°C for 15 minutes with slight shaking. Finally, absorbance was measured at 562nm in a plate reader and concentrations of samples calculated based on the BSA standard curve. HEK cell samples were diluted 1:10 before the BCA measurement. *Preparation of nuclear/cytoplasmic fractionations.* For fractionation HEK cells the NE-PER Nuclear and Cytoplasmic Extraction Reagent kit (Thermo Fisher Scientific, 78833) was used and the manufacturer’s protocol for isolation of protein fractions followed. All steps were performed at 4°C if not mentioned otherwise. Briefly, HEK cells were washed 1x with 1X PBS and detached using TrypleE for 2 min. After collection centrifuging of the cells at 500g for 3 min supernatant was discarded and 200µL CERI (containing 1X HALT protease inhibitors) added per pellet. Samples were vortexed on the lowest settings for 15s and incubated on ice for 10min. Then 11µL of CERI were added to each sample, followed by vortexing for 5s, 1min incubation on ice and another vortexing for 5s. To collect the cytoplasmic fraction samples were centrifuged at 16000g for 5min. The remaining pellet was resuspended in NER (containing 1X HALT protease inhibitors) and vortexed on the highest setting for 15 s. This step was repeated every 10min for a total duration of 40min. After the last vortex step samples were centrifuged at 16000g for 10min and supernatant (nuclear fractions) collected. Samples were immediately used for western blot procedures or stored at - 20°C. *Differential solubility assay.* Wells were washed 1x with 1XPBS and 200µL of 1% triton X-100 in lysis buffer (25mM Tris-HCl pH 7.4, 62.5mM NaCl, 1X HALT protease inhibitors, 1X Pierce NEM (Thermo Fisher Scientific, 23030) added. Cells were detached using the flat end of a syringe plunger and pipetted 15times to ensure lysis of cells. After incubation on ice for 15min samples were centrifuged for 5min at 12000rpm and supernatant (soluble fraction) was snap frozen on dry ice. The pellet (insoluble fraction) was resuspended in 100µL of 2% SDS in lysis buffer and immediately boiled at 95°C for 5min. Samples were directly used for western blot procedure or snap frozen in dry ice. *Click-iT assay.* To asses de novo protein synthesis the Click-iT AHA (L-azidohomoalanine, Invitrogen, C10102) assay was used and the manufacturers protocol followed. Briefly, 1*106 MNs were seeded in p24 well plates and methionine-free medium added for 60min. MNs were then incubated with 1X labelling reagent for 4h. After lysis of the cells samples were vortexed for 5min and after centrifugation at 13000g for 5 minutes at 4°C a BCA measurement was performed. The Click reaction was performed as indicated by the manufacturers protocol and samples immediately used for western blotting or stored at -20°C. *Western blot procedure.* All samples were run with the same protein amount and volume within one gel. Lysis buffer was used to adjust samples concentrations. Before loading samples into AnykD Criterion TGX Precast Midi or Mini Protein gels, Laemmli Buffer (BioRad, sc-40675-PR), containing 10% 2-ß-mercaptoethanol, was added to achieve a 1X concentration. Gels were run in 1x Tris-Glycine SDS Running Buffer (Thermo Fisher Scientific, LC26755) using 70V for ∼20min and 100V for ∼2hours. After running, the gel was equilibrated in 1x Tris/Glycine Transfer Buffer (BioRad, 1610734). Protein transfer was performed using pre-made TransBlot Turbo-Transfer Pack in the semi-dry Trans-Blot Turbo System (Bio-RAD, 1704150). After the transfer, the membrane was rinsed with diH2O and stained with Ponceau to check even loading. Membranes were blocked with 5% non-fat dried milk powder in 1x TBST for 1 hour at. The desired antibody was then added and incubated in PBS containing 3% BSA and 0.01% sodium azide at 4°C overnight. On the next day the membrane was washed 4 times with 1X TBST for 7min each before the secondary antibody diluted in 5% non-fat milk (PanReac AplliChem ITW, A0830) in PBS was added for 1h at RT. After another step of washing 4 times with 1X TBST for 7min each the signal was developed using western blot substrate PICO or DURA (Thermo Fisher Scientific, 34580), X-Ray Films (Fuji, 4741019289), and the Cawomat 2000 IR X-Ray developer. Densitometric analysis was performed on scanned X-Ray films using the Quantity One software (Bio-RAD).

### Microfluidic device (MFD) production & imaging procedure

MFDs were produced at the Microstructure Facility (MSF, CRTD) based on a previously published protocol (*52*). Briefly, polydimethylsiloxane (PDMS) was used to form a negative imprint of a photoresist master template that contains the structures for wells and micro-channels of the MFD. After coating of glass coverslips with polyethylene-alt-maleic anhydride PEMA, the PDMS replica was placed on the PEMA-coated coverslips and bonding induced by placing a weight on top of the PDMS replica for 2h at 120°C. The result of this bonding step is the finished MFD with PDMS wells, connected by 100 micro-channels, placed on a glass coverslip. Before the PDL coating, MFDs were treated with air plasma to hydrophilize the micro-channels (plasma time: 4min, Gas stb time: 50s, Power setpoint: 50W, Gas1(Air) setpoint 0%, Gas2 (O2) setpoint: 60%, base vacuum: 0.5mbar), for subsequent PLO coating. MFDs were incubated with a 30% PLO (Sigma Aldrich, 27378-49-0) solution in PBS, using 50µL per well, overnight at 37°C. On the following day wells were washed with 1X PBS 3times for 30 minutes each. During the last washing step MFDs were placed under UV at RT to ensure sterility. Next 25µg/mL laminin diluted in PBS were incubated in the wells overnight at 37°C. Before cells were seeded into the wells at a density of 7.5*104 wells were incubated with medium for 30min. On the day of live imaging cells were stained with LTR as explained above for 1h, by adding the dye to both compartments of the well. Imaging took place in a Zeiss Spinning Disk microscope equipped with a cage incubator, CO2 incubation and heating at 37°C. The 40x/0.95 Plan-Apochromat, Air was used for image acquisition together with the ZenBlue software (Zeiss). Videos were acquired with frames taken every 1s for a total duration of 120s. A minimum of 6 fields (each containing 8 channels) were imaged per cell line.

### Lysosomal trafficking analysis

Acquired movies were bleach corrected using the “histogram” method in FIJI (*125*). Lysosomes were tracked within the micro-channels, using the MATLAB based software “Fluorescence Image Evaluation Software for Tracking and Analysis” (FIESTA) with parameters specified in Supplemental Table 2. All automatically generated tracks were manually verified, and faulty tracks were either corrected or excluded. The tracks were then parsed into phases of positive, negative and stationary segments, using python-based segmentation algorithm. Briefly, the original data was approximated by a cubic spline (weighted least square interpolation from python package scipy.interpolate) over a three-point window (measurement interval = 1 s) to calculate the velocity at each data point. Data points were given either of two states - paused (velocity < 100 nm/s) or running (velocity ≥ 100 nm/s). To reduce noise, isolated single state points were eliminated by converting them to match their neighbouring states, after which adjacent regions of the same state were combined into segments. Runs were characterized as positive (soma to axon) or negative (axon to soma). Based on the composition of runs for each track they were classified as either anterograde (only positive runs), retrograde (only negative runs), or bi-directional (at least one positive and one negative run). If a vesicle moved less than 2.3 µm over the total 120 s imaging period, they were classified as stationary.

### Zebrafish experiments

Zebrafish were kept and bred according to standard procedures (*126*). The experimental procedures were performed on fish younger than 5dpf. The Tg(mnx1:EGFP) motor neuron reporter line (mnx1:GFP; (*127*)) was used (project license number TVV 36/2021).

*Plasmid generation for zebrafish experiments.* To generate the plasmids used in various zebrafish experimental procedures the following vectors and plasmids were used: pCS2P+ (Addgene plasmid # 17095), pminiTol2 (Addgene plasmid # 31829), pcDNA3.1-TFEBS211A- myc (Addgene plasmid # 99957), pCIP-caTFEB (Addgene plasmid # 79013) and full length human TFEB (mRNA sequence based on vector VB220110-1212ayd by VectorBuilder). The FLAG-tagged full length human TFEB and FLAG-tagged TFEB S211A was obtained by PCR (forward EcoRI 5’ – GCG GAA TTC AAC ATG GAT TAT AAA GAT GAT GAT GAT AAA CCA GGA CCA ATG GCG TCA CGC ATA GGG -3’ and reverse XbaI 5’ - GCG TCT AGA TTA CAG CAC ATC GCC CTC C -3’) and subcloned into the empty pCS2P+ vector. Next, the FLAG-TFEB-SV40pA and FLAG-TFEB S211A-SV40pA fragments were amplified by PCR from pCS2P+-FLAG-TFEB and pCS2P+-FLAG-TFEB S211A plasmids (forward EcoRI 5’ – GCG GAA TTC AAC ATG GAT TAT AAA GAT GAT GAT GAT AAA CCA GGA CCA ATG GCG TCA CGC ATA GGG -3’ and reverse EcoRV 5’ – GCG GAT ATC AAA AAA CCT CCC ACA C - 3’), and subcloned into the EcoRI-EcoRV-digested pminiTol2-mnx1 vector to generate the final plasmids (pminiTol2 -mnx1:FLAG-TFEB fl and pminiTol2-mnx1:FLAG-TFEB S211A). The truncated TFEB fragment was generated by removing the first 400 bp from the N-terminal position by PCR (forward EcoRI 5’ – GCG GAA TTC AAC ATG GAT TAT AAA GAT GAT GAT GAT AAA CCA GGA CCA ATG CTG CAC ATT GGC TCC AAC – 3’ and reverse XbaI 5’ - GCG TCT AGA TTA CAG CAC ATC GCC CTC C -3’) and subcloned into the pCS2P+ vector to generate the pCS2P+-FLAG-trTFEB plasmid. The FLAG-trTFEB-SV40pA fragment was amplified by PCR (forward EcoRI 5’ – GCG GAA TTC AAC ATG GAT TAT AAA GAT GAT GAT GAT AAA CCA GGA CCA ATG CTG CAC ATT GGC TCC AAC – 3’ and reverse EcoRV 5’ – GCG GAT ATC AAA AAA CCT CCC ACA C - 3’), and subcloned into the EcoRI-EcoRV-digested pminiTol2-mnx1 vector to generate the pminiTol2-mnx1:FLAG-trTFEB plasmid. TFEB without the DNA binding domain was generated by PCR using the full length TFEB as template (forward EcoRI 5’ – GCG GAA TTC AAC ATG GAT TAT AAA GAT GAT GAT GAT AAA CCA GGA CCA ATG GCG TCA CGC ATA GGG – 3’ and reverse XbaI 5’ - GCG TCT AGA TTA CCG CTC CTT GGC CAG G – 3’) and ligated into the EcoRI-XbaI-digested pCS2P+ vector. The FLAG-TFEB w/o DBD-SV40p2A fragment was subcloned with EcoRI and EcoRV into the pminiTol2-mnx1 vector (forward EcoRI 5’ – GCG GAA TTC AAC ATG GAT TAT AAA GAT GAT GAT GAT AAA CCA GGA CCA ATG GCG TCA CGC ATA GGG – 3’ and reverse EcoRV 5’ – GCG GAT ATC AAA AAA CCT CCC ACA C - 3’). The *mnx1* promoter fragment was amplified by PCR from the pminiTol2-mnx1 vector (*128*) and inserted into an empty pminiTol2 vector with BglII and NotI to generate the pminiTol2-mnx1 vector (forward BglII 5’ – GCG AGA TCT CCA TTT AAA TTA GCC TGG CAT CTG GAC – 3’ and reverse NotI 5’ – GCG GCC GCT CTG GCC CAC CTC ACA AAC AG – 3’). All generated plasmids were fully or partially sequenced.

*Morpholino and DNA plasmid injections.* Tg(mnx1:EGFP) embryos were injected with a mixture containing Tol2 transposase mRNA (40ng/ml, pT3TS-Tol2, Addgene plasmid # 31831), DNA plasmid containing one of the TFEB constructs (5ng/ml), and SMN-MO (McWhorter et al., 2003) (3 ug/ul, 5’ – CGACATCTTCTGCACCATTGGC – 3’, Gene Tools, LLC, USA) at the single-cell stage (final volume: 1 nl). mRNA coding for Tol2 transposase was generated by linearizing the pT3TS-Tol2 (plasmid Addgene #31831) with XbaI, followed by mRNA synthesis (Ambion™ mMESSAGE mMACHINE T3 transcription kit, AM1348, Thermo Fisher Scientific) and purification (RNeasy Mini Kit, Roche).

*CaP motor axon morphology analysis and motor neurons counts.* Motor axon morphology was evaluated at 26-28 hpf. Embryos were fixed in 4% PFA at room temperature for 1-2 hours, followed by mounting in 70% glycerol and image acquisition using the Apotome unit AxioCam MRm (60N-C) equipped with a LD LC1 Plan-Achromat 20x-objective (NA=0.8). Images were acquired using the Zeiss Zen Black Software and an AxioCam MRm (60N-C). Maximum intensity projections were generated in Fiji software. Calculation of the % of aberrant CaP MNs was based on a previously published protocol (*129*). Briefly, CaP motor axons at the developmental stage of 26-28 hpf that show more than 2 branches, are truncated or missing are categorized as aberrant. Between six and seven axons per hemisegment (caudally from somite 7) were scored. Both sides of the embryos were analyzed, with a total of 12-14 axons/embryo being scored. Two independent observers analyzed the morphology of the CaP axons. During all analyses the observers were blinded to the treatments (randomization and encryption of file was done using the Blind analysis Tool v1.0 plugin in Fiji software). For the analysis of the number of motor neurons a region of interest (ROI) was defined at the level of the spinal cord with dimensions of: width = 200 um, height = 50 um and depth = 10 um. The number of cells was done manually using the Zen Blue software. The observer was blinded to the experimental conditions using the Blind analysis Tool v1.0 plugin in Fiji software.

*Zebrafish whole-mount immunohistochemistry.* For immunostaining of the FLAG-tagged TFEB constructs, embryos (26-28 hpf) were dechorionated and fixed in 4% PFA for 45 minutes at RT. Next, embryos were blocked in blocking buffer (1% DMSO, 1% BSA, 0.5% Triton X, 1x PBS with 5% donkey serum) for 1-2h, followed by overnight incubation in 300 μL blocking solution containing anti-FLAG M2 (1:100, Sigma, F1804) and anti-GFP antibodies (1:300, abcam, ab13970). After washing, embryos were incubated in donkey anti-mouse secondary antibody labelled with Alexa Fluor 546 (1:200; Jackson ImmunoResearch) and donkey anti-rabbit secondary antibody with Alexa Flour 488 (1:200, Jackson ImmunoResearch). Microscopical analysis was performed in 70% glycerol on micro slides using the Apotome unit (AxioCam MRm (60N-C) with a LD LC1 Plan-Achromat 20x-objective (NA=0.8). Images were acquired using the Zeiss Zen Black Software and an AxioCam MRm (60N-C).

### Antibodies used for human cells

For cell imaging and Western blot experiments, the following antibodies were used: mouse SMN (BD Bioscience 610646), mouse actin (Santa Cruz, A1416), rabbit BRN2 (Cell Signaling, 12137), rabbit ISL1 (Abcam, ab109517), mouse ISL1/2 (Hybridoma Bank, 39.4D5-c), chicken MAP2 (NB300-213, Novus Biologicals), β-III Tubulin (Merck Millipore, 9354), rabbit CTSB (Cell Signaling, 31718), mouse Lamp1 (Hybridoma Bank, H4A3-s), rabbit lamin B1 (Abcam, ab16048), mouse p62 (Abcam, ab56416), rabbit p-S6K1 (Thr389, Cell Signaling Technology, 9234S), rabbit S6K1 (Cell Signaling Technology, 2708S), rabbit p-S6 (Ser235/236, Cell Signaling Technology, 2211s), rabbit S6 (Cell Signaling Technology, 2217S), rabbit TFEB (Cell Signaling, 4240S), rabbit vimentin (Cell Signaling, 5741S), rabbit ubiquitin (Santa Cruz Biotechnology, SC-9133). Secondary antibodies were all purchased from Thermo Fisher Scientific.

### Statistical Analysis

Statistical significance was determined using GraphPad Prism 10.3.1 (Graphpad Software, Inc.). To test the Gaussian distribution of residuals Shapiro-Wilk test was performed. Paired two-tailed t-Test was used for datasets composed of one variable (e.g. genotype) and two cell lines. If no Gaussian distribution of the residuals, nonparametric Wilcoxon matched-pairs signed rank test was performed. One-way ANOVA was performed for datasets composed of only one variable and more than 2 groups and Two-way ANOVA was chosen for datasets composed of two variables. Multiple comparison correction was applied to all ANOVA tests and is specified in each Figure legend. “N” indicates the number of independent biological experiments; “n” indicates the number of technical replicates (i.e. individual wells per experiment). A confidence interval of 95% was used for all comparisons. Graphs indicate mean+SEM unless otherwise indicated. *p<0.05, **p<0.01, ***p<0.005, ***p<0.001, ****p<0.0001. “ns” means not significant.

## List of Supplementary Materials

Fig. S1 to S8

Table S1 and Table S2

## Acknowledgments

We deeply thank Brunhilde Wirth and Michele Marass for their critical and constructive feedback on the manuscript. We would like to thank Silke White (DZNE Imaging Facility) for their support and training on multiple microscopes and Tomasz Basiewicz (CRTD Microstructure Facility) for the training and support on MFD usage and Jessica Bilstein for the training and experimental help for live trafficking experiments.

## Author Contributions

Conceptualization: NRM

Methodology: IR, AC, ZD, TS, AMO, SU, RG, HG, TG, NRM

Investigation: IR, TG, NRM

Funding acquisition: CB, SD, NRM

Project administration: NRM

Supervision: CB, BF, SD, TG, NRM

Writing – original draft: IR, NRM

Writing – review & editing: NRM

## Data and materials availability

The use of the hiPSCs (BJ WT, SMA and isogenic corrected lines) is restricted by a material transfer agreement (MTAs).

## SUPPLEMENTARY MATERIAL

**Supplemental Figure 1.**
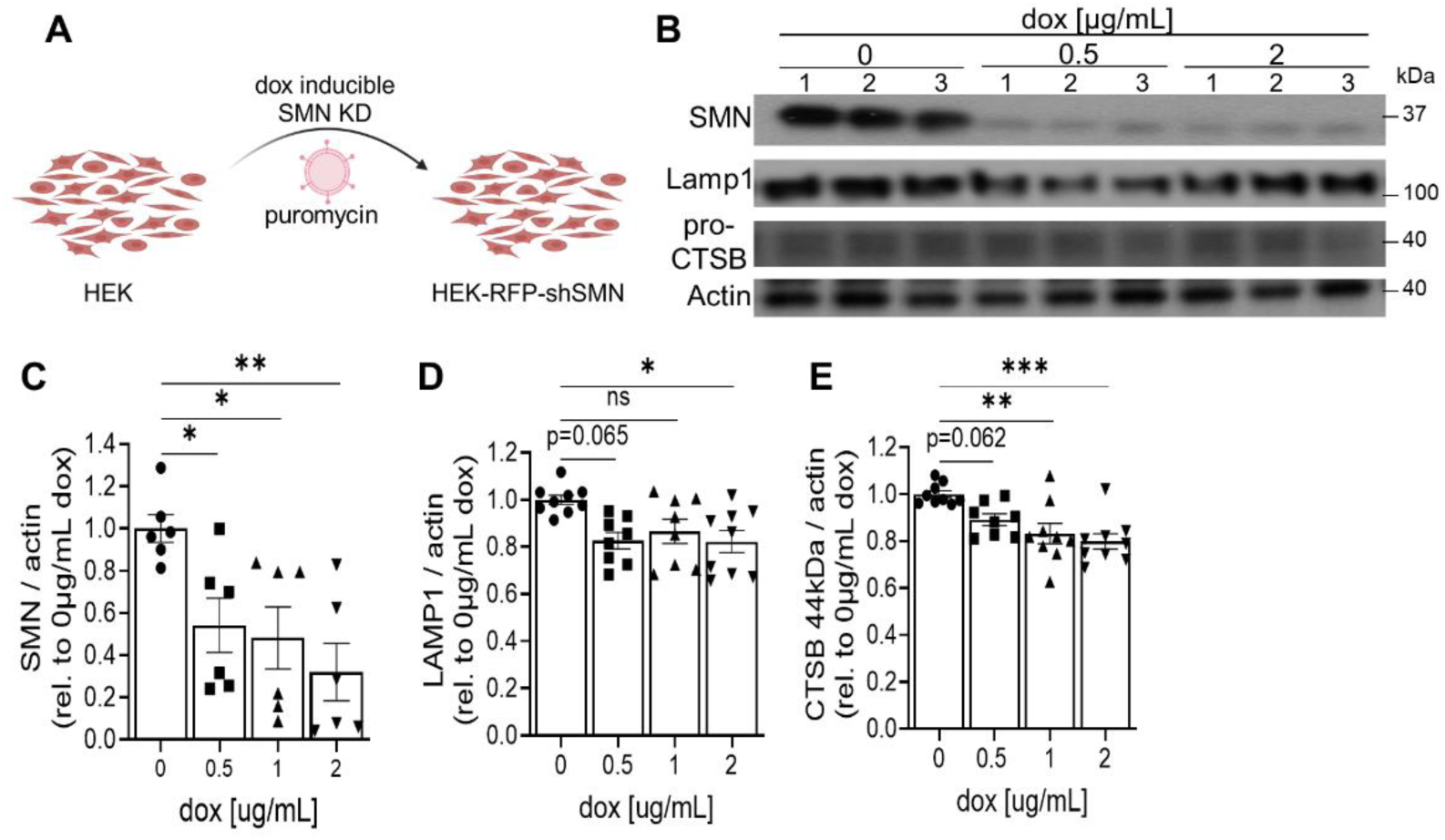
SMN reduction leads to decreased levels of lysosomal associated proteins. (**A**) Schematic representation of the generation of a stable dox-inducible SMN KD HEK line (HEK-RFP-shSMN). (**B**) Representative immunoblot of protein lysates from HEK-RFP-shSMN cells treated with the indicated dox concentrations for 72 hours and corresponding quantification of (**C**) SMN, (**D**) LAMP1 and (**E**) CTSB protein levels (One-way ANOVA with Bonferroni’s multiple comparison test, N= 2-4 independent experiments. Mean ± SEM is shown).

**Supplemental Figure 2.**
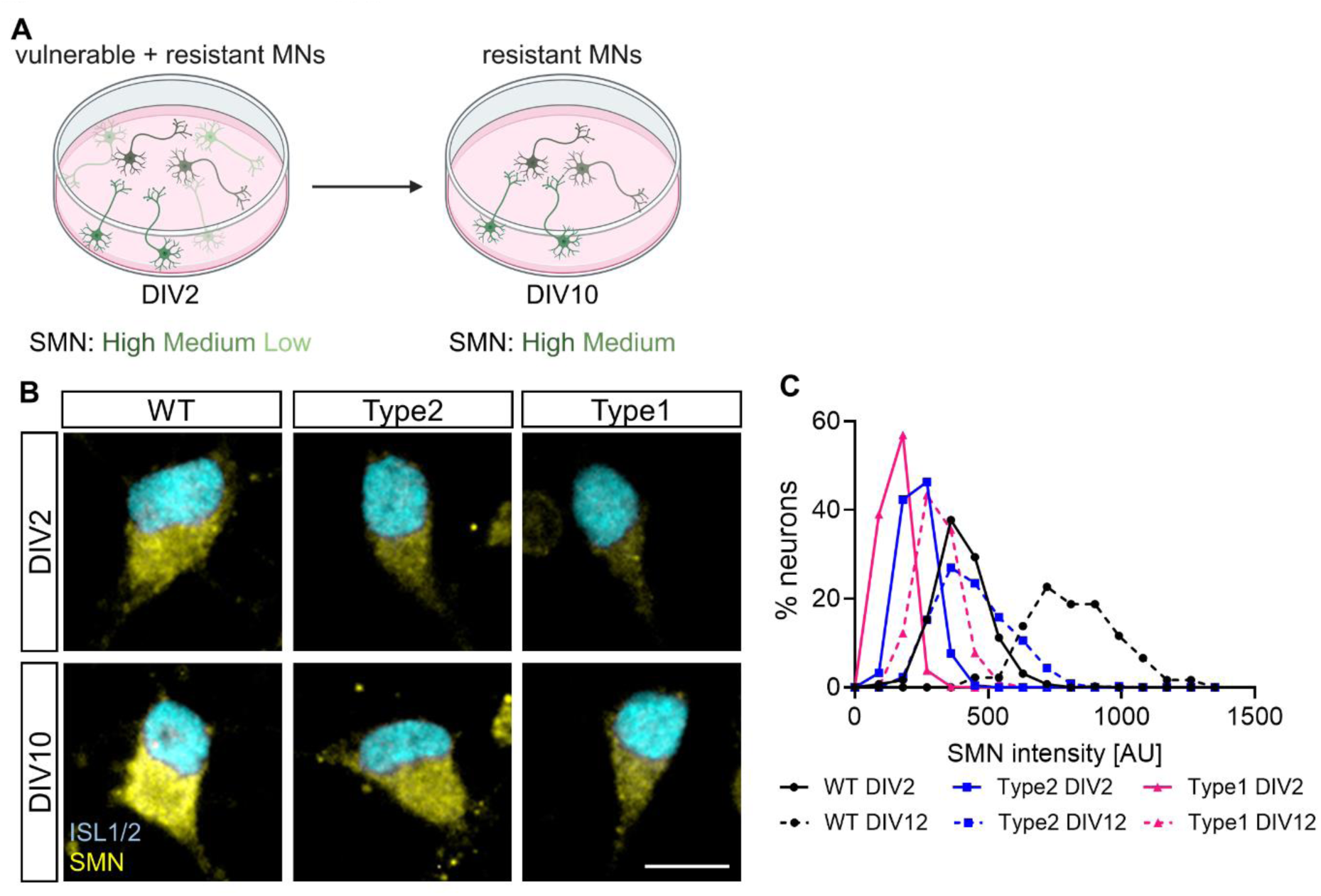
The resilient MNs that survive longer in culture harbour higher SMN protein levels than the ones present in earlier cultures. (**A**) Schematic representation of differential survival of vulnerable and resistant MN populations according to their intrinsic SMN protein levels. (**B**) Representative images of DIV2 and DIV10 WT, SMA type 2 and type 1 MNs immunostained against SMN (orange) and ISL1/2 (cyan). Scale bar, 10µm. (**C**) Representative histogram analysis showing the percentage of WT, SMA type 2 and type 1 MNs falling into each SMN intensity bin after 2 and 12 days in vitro.

**Supplemental Figure 3.**
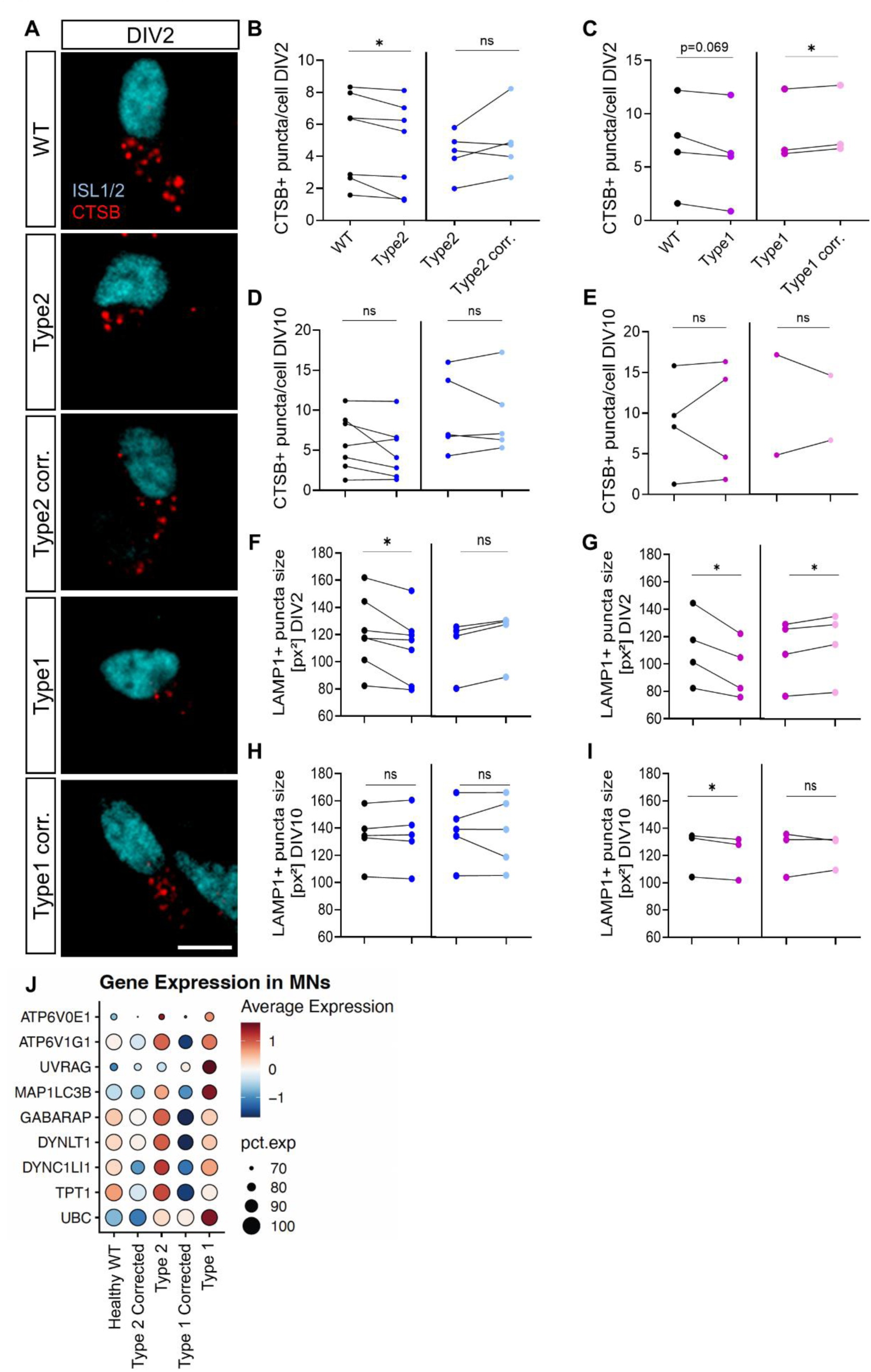
SMA MNs showed decreased number of lysosomes and lysosomal size in early but not in late cultures. (**A**) Representative images from DIV2 MNs derived from WT, SMA type 2 and type 1 and their respective isogenic control hiPSCs immunostained against CTSB (red) and ISL1/2 (blue). Scale bar, 10µm. Quantification of the number of CTSB^+^ puncta per ISL1/2 ^+^ MN from SMA type 2 vs WT and isogenic control (**B**) and SMA type 1 vs WT and isogenic control (**C**) at DIV2 and DIV 10 (**D-E**). (**F**) Quantification of the number of LAMP1^+^ puncta size in ISL1^+^ DIV2 MNs from SMA type 2 vs WT and isogenic control and (**G**) SMA type 1 vs WT and isogenic control and similar quantifications on DIV10 MNs (**H-I**) (For all panels, 2-tailed Paired t-test, each circle represents an independent experiment, each representing the mean of n= 3 technical replicates, samples from the same experiment are connected by a line. Mean ± SEM is shown). (**J**) Dot plot showing expression levels of autophagy/lysosome related genes from the MN subpopulations of day 20 neuromuscular organoids subjected to scRNAseq (19).

**Supplemental Figure 4.**
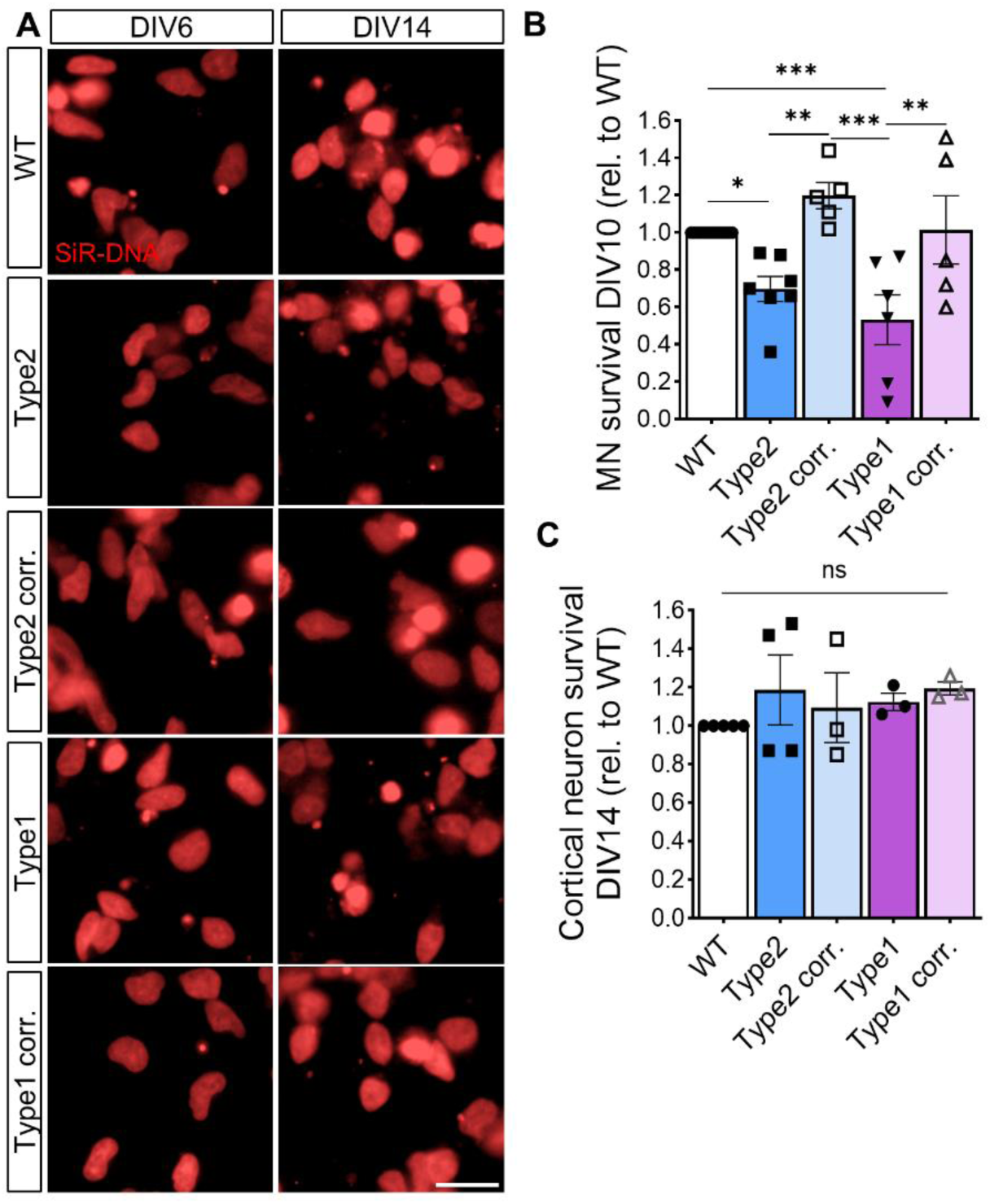
SMN protein is essential for the survival of MNs, but not for CxNs. (**A**) Representative images of cortical neurons (CxNs) 6 and 14 days after LV-mediated *NGN2* overexpression labelled with the live dye SiR-DNA. Scale bar, 10µm. (**B**) Quantification of the percentage of SiR-DNA-labeled MNs that survived after 10 days in culture relative to the ones detected 2 days after plating. (**C**) Quantification of the percentage of SiR-DNA-labeled CxNs that survived after 14 days in culture relative to the ones observed 6 days after plating and LV-*NGN2* overexpression (One-way ANOVA with Bonferroni’s multiple comparison correction. Mean ± SEM is shown).

**Supplemental Figure 5.**
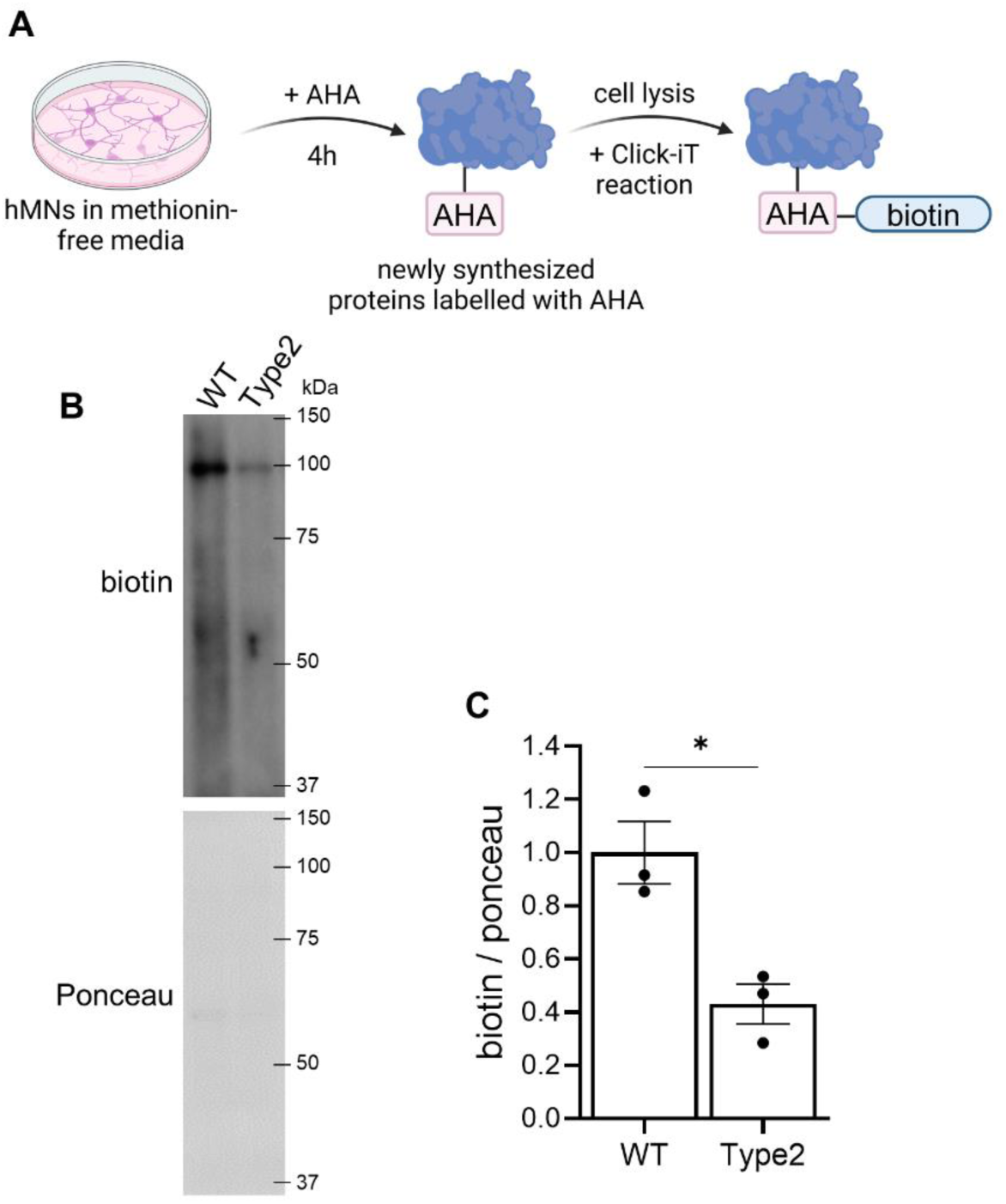
The protein aggregation feature observed in SMA MNs is not due to an enhanced protein synthesis. (**A**) Schematic of the Click-iT assay followed to label newly synthesized proteins. (**B**) Representative immunoblot of protein lysates from WT and SMA type 1 MNs subjected to protein synthesis Click-iT assay showing biotin levels and (**C**) quantification of newly synthesized proteins (2-tailed Paired t-test, each circle represents an independent experiment. Mean ± SEM is shown).

**Supplemental Figure 6.**
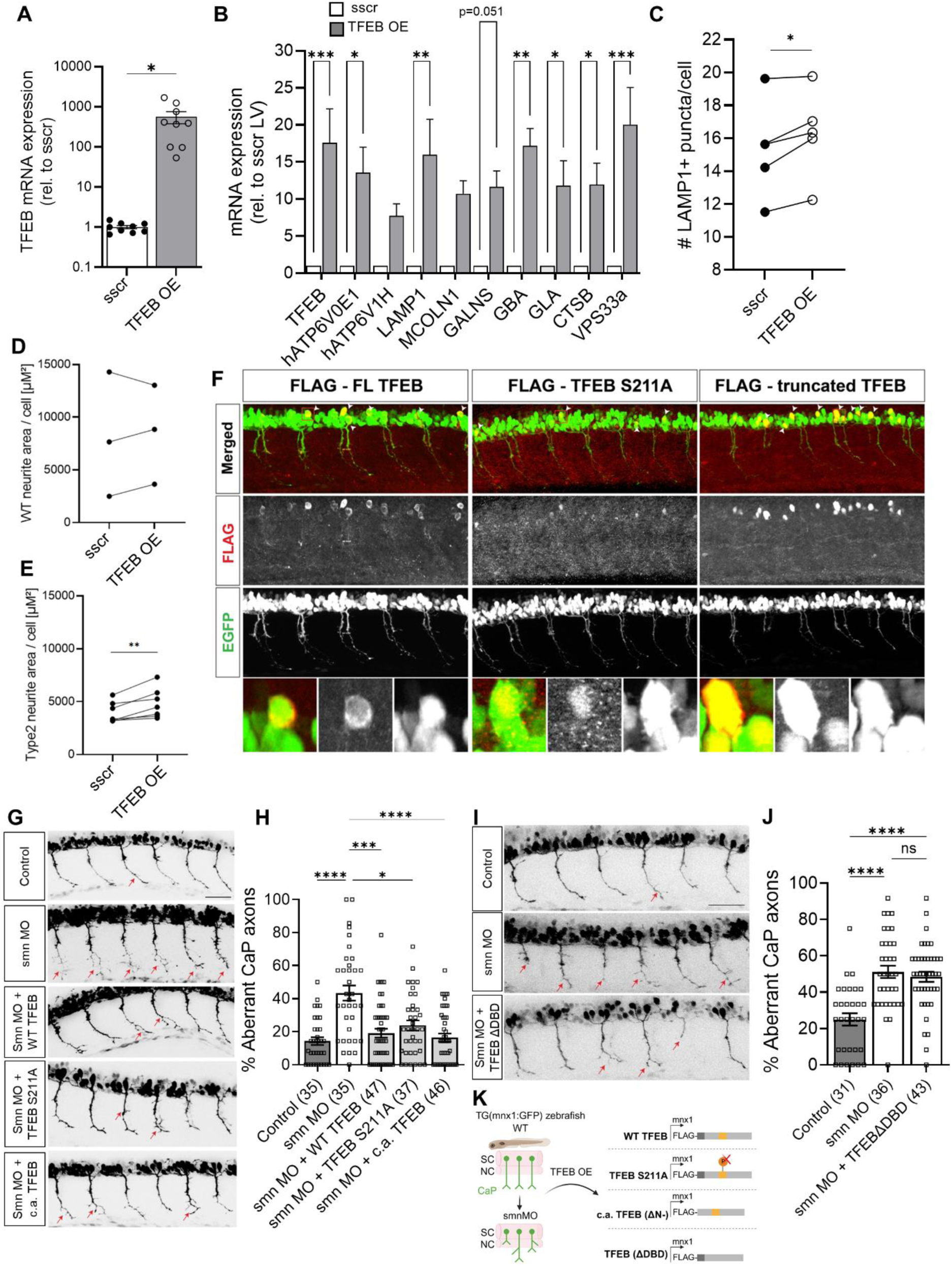
TFEB overexpression increases CLEAR gene expression levels and the number of lysosomes. (**A**) Quantification of TFEB mRNA levels from DIV10 WT hiPSC-derived MNs after transduction with lentivirus (LV) carrying sscr control or TFEB for 9 days (N = 2 independent experiments, each composed by n =3 technical replicates). (**B**) mRNA expression levels of CLEAR genes measured by qPCR from HEK293T cells transduced with either sscr control or TFEB-overexpression LV for 4 days (two-way ANOVA with matched values and Bonferroni’s multiple comparison test, N= 3 independent experiments, each composed by n= 2 technical replicates). (**C**) Quantification of the number of LAMP1^+^ puncta per MN at DIV10 after sscr or TFEB-overexpression (2-tailed Paired t-test, each point represents an independent experiment composed of n= 3 technical replicates. Samples from the same experiment are connected by a line). Quantification of neurite length per MN after sscr or TFEB-overexpression in healthy control (**D**) and SMA type 2 MNs (**E**) (2-tailed Paired t-test, each point represents an independent experiment composed of n= 3 technical replicates. Samples from the same experiment are connected by a line). (**F**) Representative immunostaining against FLAG showing spinal motor axons of a zebrafish SMA model upon injection of smn morpholino (MO) + WT TFEB; smn MO + TFEB S221A or smn MO + truncated TFEB 28 hours after fertilization. (**G, I**) Representative micrographs of spinal motor axons from a zebrafish SMA model 28 hours after fertilization and morpholino +/- TFEB construct injection. Scale bar, 50μm. (**H, J**) Quantification of the percentage of spinal MNs with aberrant axons for the indicated experimental conditions (Kruskal-Wallis test with Dunn’s post hoc test). Mean ± SEM is shown. (**K**) Schematic representation of TFEB constructs used for over expression in a Zebrafish SMA model (Biorender).

**Supplemental Figure 7.**
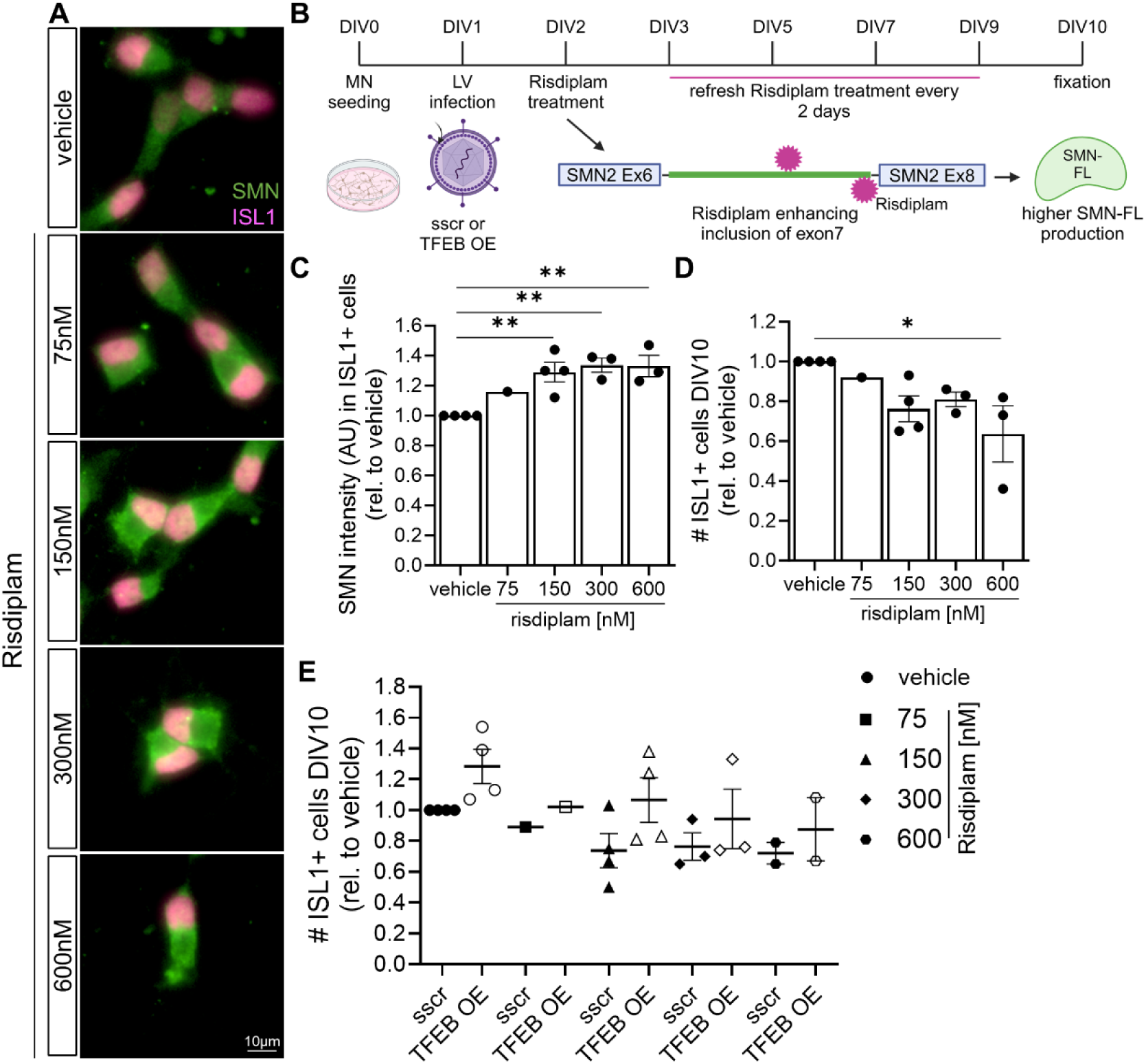
Combinatorial treatment to enhance TFEB levels and correct *SMN2* defective splicing shows no additional effect on MN survival. (**A**) Representative images of DIV10 SMA type 2 hiPSC-derived MNs treated with risdiplam or vehicle for 9 days and immunostaining against SMN (green) and ISL1 (magenta). Scale bar, 10µm. (**B**) Schematic experimental design for the combinatorial treatment (TFEB overexpression and risdiplam) on hiPSC-derived MNs. (**C**) Single cell quantification of SMN intensity in ISL1^+^ MNs after treatment with the indicated concentrations of risdiplam or vehicle for 9 days. (**D**) Quantification of the number of DIV10 ISL1^+^ SMA type 2 MNs after risdiplam treatment (One-way ANOVA with Bonferroni’s multiple comparison correction, each symbol represents an independent experiment, each comprised of n= 3 technical replicates). (**E**) Quantification of the survival of MNs after transduction with the LVs carrying either sscr or TFEB and treated with risdiplam or vehicle for 9 days (results are expressed relative to “sscr LV + vehicle” condition) (Two-way ANOVA, each symbol represents an independent experiment, each comprised of n= 3 technical replicates). Mean ± SEM is shown.

**Supplemental Figure 8.**
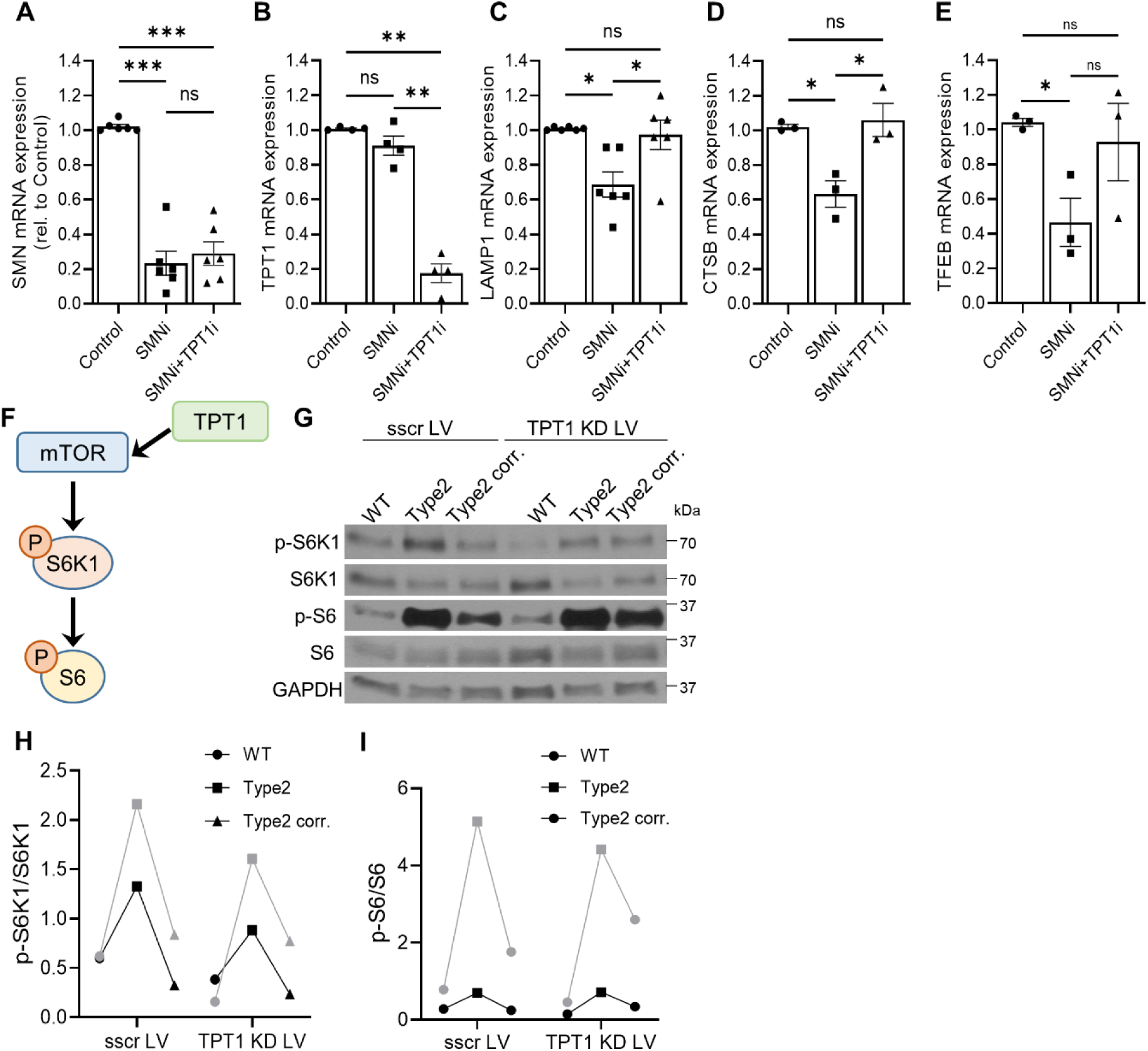
TPT1 levels are increased in SMA MNs and its knockdown ameliorates mTOR overactivation. Quantification of (**A**) *SMN*, (**B**) *TPT1*, (**C**) *LAMP1*, (**D**) *CTSB* and (**E**) *TFEB* mRNA expression levels measured by qPCR from HEK293T cells treated with control RNAi or RNAi against *SMN1* or *SMN1*+*TPT1* (One-way ANOVA with Tukey multiple comparison test, each point represents an independent experiment. Mean ± SEM is shown). (**F**) Simplified scheme showing TPT1-mTOR signalling pathway. (**G**) Representative immunoblot of protein lysates from WT and SMA type 2 MNs treated for 7 days with lentivirus carrying sscr control or shRNA against *TPT1* and quantification of (**H**) phosphorylated S6K1 and (**I**) phosphorylated S6 over total protein levels for each of the MN conditions (N = 2 independent experiments, each representing the mean of n= 3 technical replicates). Mean ± SD is shown.

## Supplemental Tables

**Supplemental Table 1:**
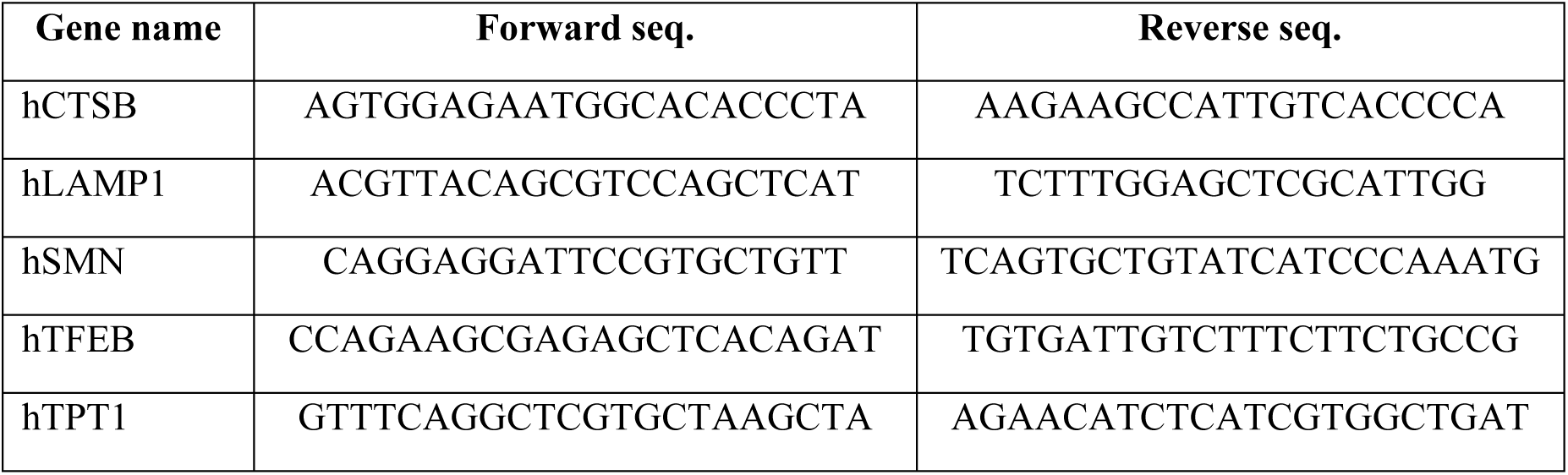
qPCR primers used.

**Supplemental Table 2:** Parameters used for image analysis in the FIESTA software.

| Parameter | Value | Parameter | Value |
| --- | --- | --- | --- |
| max. velocity | 2.4μm/s | pixel size | 114nm |
| time difference | 1000ms | only molecules | true |
| height | 2 | Fit(CoD) | 0 |
| FWHM(Est.) | 500nm | border margin | 1 pixel |
| verification steps | 2 | max. angle | 45° |
| min. length | 4 frames | max. break | 4 frames |
| weights: position,<br>direction, speed | 40%, 40%, 20% | Model | Symmetric 2D-Gaussian |

